# Variant Surface Protein GP60 Contributes to Host Infectivity of *Cryptosporidium parvum*

**DOI:** 10.1101/2024.02.04.578776

**Authors:** Muxiao Li, Fuxian Yang, Tianyi Hou, Xiaoqing Gong, Na Li, L. David Sibley, Yaoyu Feng, Lihua Xiao, Yaqiong Guo

## Abstract

Biological studies of the determinants of *Cryptosporidium* infectivity and virulence are lacking despite the fact that cryptosporidiosis is a major public health problem. Here, we used advanced genetic tools to investigate the processing, fate, and function of the 60-kDa glycoprotein (GP60), an immunodominant variable antigen associated with protection against reinfection. Endogenous gene tagging revealed that GP60 is highly expressed in sporozoites, merozoites and male gametes, suggesting that it may be involved in both invasion and sexual replication. GP60 is translocated to the parasite membrane and cleaved at the furin cleavage site into GP40 and GP15. During invasion, GP40 translocates to the apical end of the zoites and remains detectable at the parasite-host interface. Although GP60 is dispensable, both gene deletion and replacement reduce parasite growth and severity of infection. Depletion of its structural domains, GP40, or GP15 individually affects GP60 translocation but has less effect on its function. These findings suggest that the GP60 protein contributes to host infectivity likely through its multiple functions in *C. parvum*-host interactions. They further our understanding of the pathogenesis of cryptosporidiosis.

## Introduction

Cryptosporidiosis is a major public health problem in both developed and developing countries (1). It causes moderate to severe diarrhea and associated stunting and mortality in children in low- and middle-income countries and large outbreaks of enteric disease in the general population in high-income countries (2, 3). Treatment for the disease is limited, with only one partially effective drug, nitazoxanide, approved by the US Food and Drug Administration, and no vaccines against the causative *Cryptosporidium* pathogen (4). This is largely due to a lack of in-depth understanding of the biological factors that control infectivity and virulence (5).

The 60-kDa glycoprotein (GP60) is one of the earliest molecules identified to be involved in *Cryptosporidium*-host interactions and possibly virulence. The gene encoding the protein was identified independently by four research groups over 20 years ago through screening expression libraries and N-terminal sequencing of monoclonal antibody identified proteins (6–9). Bioinformatic analysis suggests that the protein is a secretory protein with an N-terminal signal peptide and a C-terminal glycophosphatidylinositol (GPI) anchor. The results of these studies suggest that the protein is likely a precursor with a cleavage site that proteolytically processes the protein into two smaller mature fragments, GP40 and GP15. A putative furin cleavage site was soon identified in the GP60 sequences, and is used by a subtilisin-like serine protease, *Cp*SUB1, to cleave GP60 into GP40 and GP15 (10, 11). The GP15 fragment of the protein was identified as an immunodominant antigen and one of the seven *Cryptosporidium* proteins associated with protection against reinfection in children (12). As an immunodominant antigen under selection pressure, GP60 is a variant protein with extensive sequence polymorphism (6). As a result, the *GP60* gene is the most widely used marker for subtyping human pathogenic *C. parvum* and *C. hominis* (13). Among the major *GP60* subtype families of *C. parvum*, the IIa, IIc, and IId subtypes differ significantly in host preference (14). Similarly, some *GP60* subtypes of *C. hominis* are more virulent than others (15).

Despite its potential importance in invasion and virulence, research into the function and action of GP60 in *Cryptosporidium*-host interactions is hampered by the mucin-like nature of both GP40 and GP15 with heavy O-linked glycosylation and the presence of a GPI anchor in GP15. As a result, recombinant GP60 produced by the prokaryotic expression system lacks some characteristic features of the native GP60 protein, making it difficult to study its biological processing and functions (16). Therefore, although the *GP60* gene is the most widely studied *Cryptosporidium* gene, only a handful of studies have been conducted on the biological properties of the GP60 protein. Even the subcellular location of GP60 is controversial. Some believe that GP60 is a surface protein attached to sporozoites through the GPI anchor (6, 17), another group suggests that GP60 is associated with the parasitophorous vacuole membrane (PVM) (18), while others have used GP60 as a marker for micronemal proteins (19).

In the present study, we have performed the first biological characterization of the GP60 protein using genetic manipulation tools. Endogenous gene tagging, depletion, and replacement were performed using the newly established CRISPR/Cas9 technology. The results of the study indicate that the multifunctional GP60 is translocated through its unique structural domains to different organelles during *C. parvum* invasion and development, and that both GP40 and GP15 are required for GP60-mediated host infectivity.

## Results

### GP60 is highly expressed *in vivo*

The results of the whole transcriptome analysis indicated that the *GP60* gene was the most highly expressed gene in both the ileum and colon of IFN-γ knockout (GKO) mice infected with *C. parvum* (Fig. 1*A*). The high expression of GP60 protein was confirmed by immunofluorescence analysis of tissue sections from infected mice using a mAb against the GP40 fragment of the protein, which showed the presence of many green fluorescence-stained parasites at the brush border of the ileum and colon. Some developmental stages were also seen deep in the intestinal tissue on the surface of the crypt of Lieberkuhn (Fig. 1*B*). The presence of high levels of GP60 expression at both the brush border and crypt of Lieberkuhn was further confirmed by immunohistochemical staining of tissue sections of the ileum and colon from infected animals using the same mAb (Fig. 1*C*).

**Figure 1.**
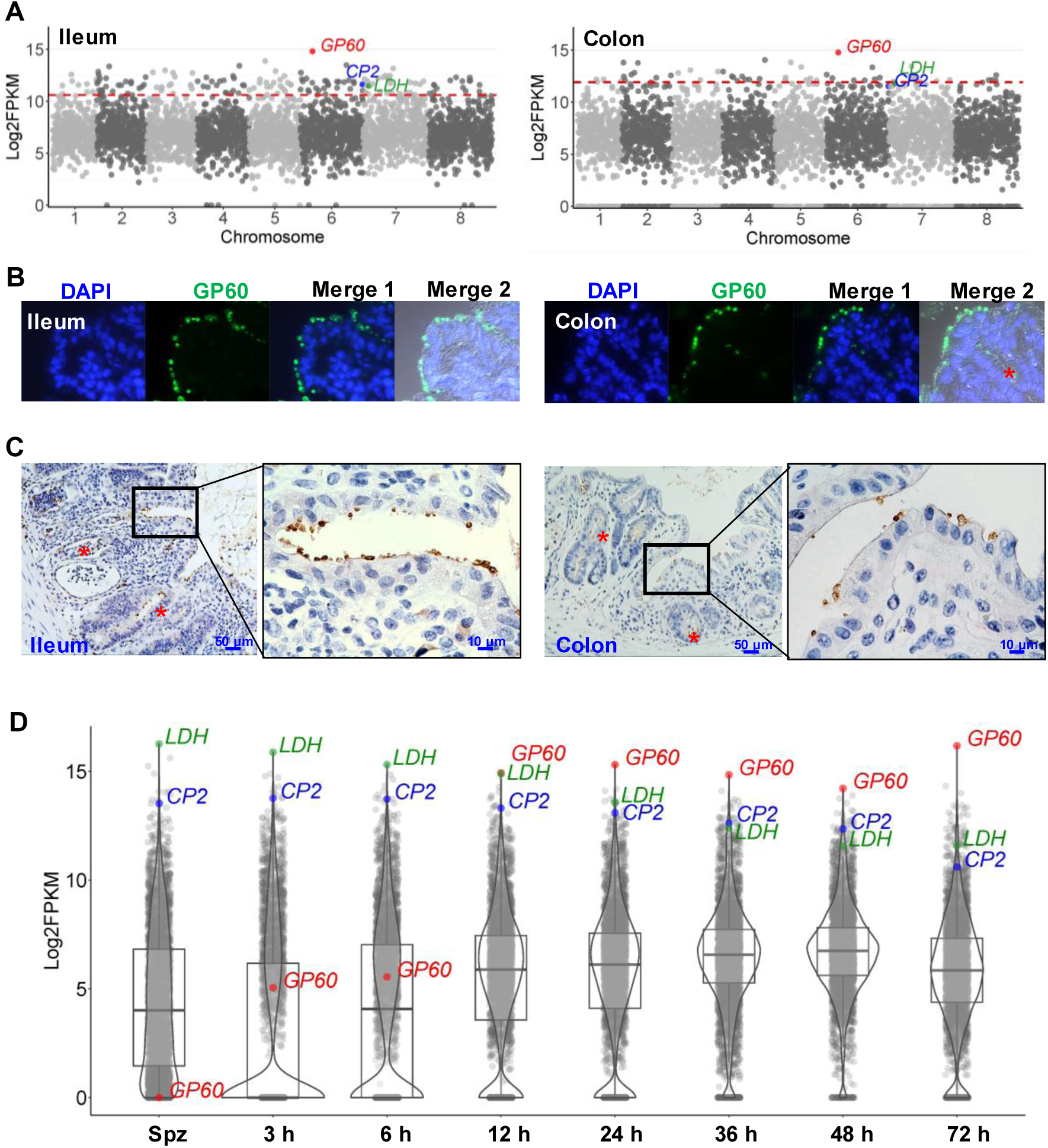
GP60 is highly expressed *in vitro* and *in vivo*. **A.** Manhattan plots showing the relative expression of all *Cryptosporidium* genes (by chromosome) in the ileum and colon of IFN-γ knockout (GKO) C57BL/6 mice infected with IIdA20G1-HLJ isolate of *Cryptosporidium parvum*, as indicating by the FPKM values in RNA-seq analysis of the transcriptome. Each dot represents the expression level of one gene, with the expression of the *GP60* gene and two genes (*CP2* and *LDH*) known to have high expression being indicated (n = 4). **B.** Expression of GP60 in ileum and colon sections as detected by immunofluorescence analysis using a mAb against recombinant GP60. *GP60* expression in parasites within the crypt of Lieberkuhn is indicated by an asterisk symbol. **C.** Expression of GP60 in ileum and colon sections as detected by immunohistochemical analysis using the *GP60* mAb. GP60 expression in parasites within the crypt of Lieberkuhn is indicated by an asterisk symbol. **D.** Violin plots showing the relative expression of all *Cryptosporidium* genes in sporozoites and developing stages in HCT-8 cells after infection with IIdA20G1-HLJ for various time. Each dot represents one gene, with the expression of *GP60* gene in comparison with the *CP2* and *LDH* genes being indicated (n = 4).

### GP60 is a surface protein expressed in invasion stages as well as in male gametes

Immunofluorescence analysis of developing stages in HCT-8 cell cultures showed high reactivity of the mAb against GP40 with free sporozoites, developing trophozoites, meronts and male gamonts (Fig. 2*A*). Immunoelectron microscopy (IEM) of parasites in intestinal tissues of infected mice revealed the presence of gold particles on the surface of late trophozoites, merozoites within meronts, and gametes within male gamonts (Fig. 2*B*). In contrast, other developmental stages such as early trophozoites and female gametes showed lower reactivity to the mAb. In mature oocysts *in vivo*, GP40 expression was mainly on the surface of sporozoites (Fig. 2*C*).

**Figure 2.**
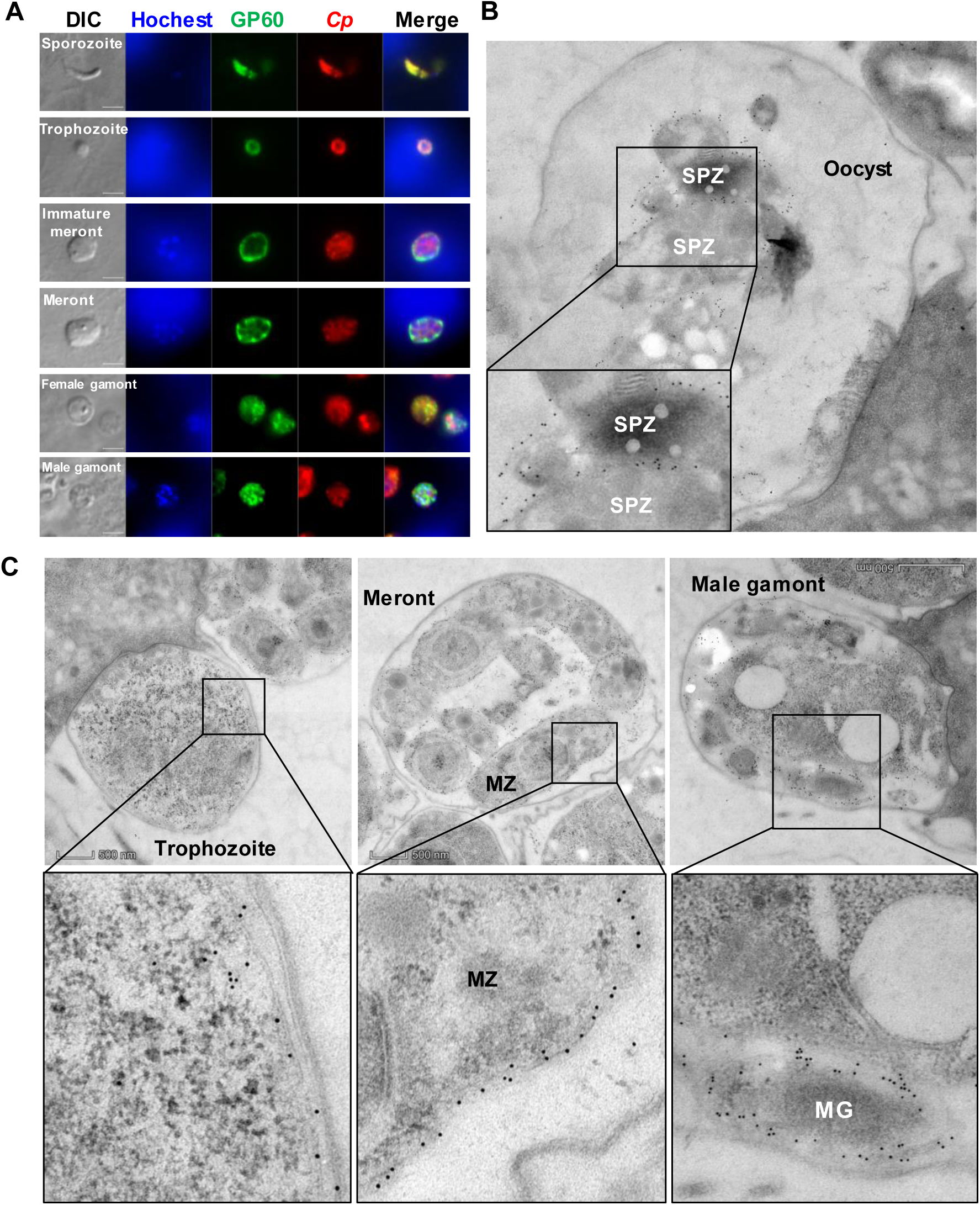
GP60 is a surface protein highly expressed in invasion stages as well male gametes. **A.** Expression of GP60 in developing stages of *Cryptosporidium parvum* cultured in HCT-8 cells, as indicated by immunofluorescence analysis using a mAb against recombinant GP60. A polyclonal antibody against sporozoites (Sporo-Glo) is used as the control (Anti-CP). **B.** Expression of GP60 in developing stages of *C. parvum* in the ileum of IFN-γ knockout (GKO) C57BL/6 mice, as indicated by immunoelectron microscopy using the mAb against GP60. Gold particles are mainly detected on the surface of late trophozoites, merozoites (MZ) within meronts, and gametes (MG) within the male gamonts. **C.** Expression of GP60 in an oocyst within the parasitophorous vacuole in the ileum of mice as indicated by immunoelectron microscopy using the GP60 mAb. Gold particles are restricted to the surface of sporozoites (SPZ).

### Endogenous gene tagging supports the membrane location GP60 protein expression

To study the dynamics of the GP60 protein and its cleavage products in the developmental stages of *C. parvum*, we performed endogenous tagging of the *GP60* gene using CRISPR/Cas9 to add a 3× HA tag and a FLAG tag in the gene after the sequence encoding the signal peptide (between amino acids 33 and 34) and the furin cleavage site (between amino acids 212 and 215), respectively, producing the expected GP40 product with an N-terminal HA tag and GP15 product with an N-terminal FLAG tag. Sequences of the neomycin resistance gene (Neo) and Nanoluciferase (Nluc) were also included for selection of the tagged line and monitoring of parasite load in mice infected with the transfected *C. parvum* (Fig. 3*A*).

**Figure 3.**
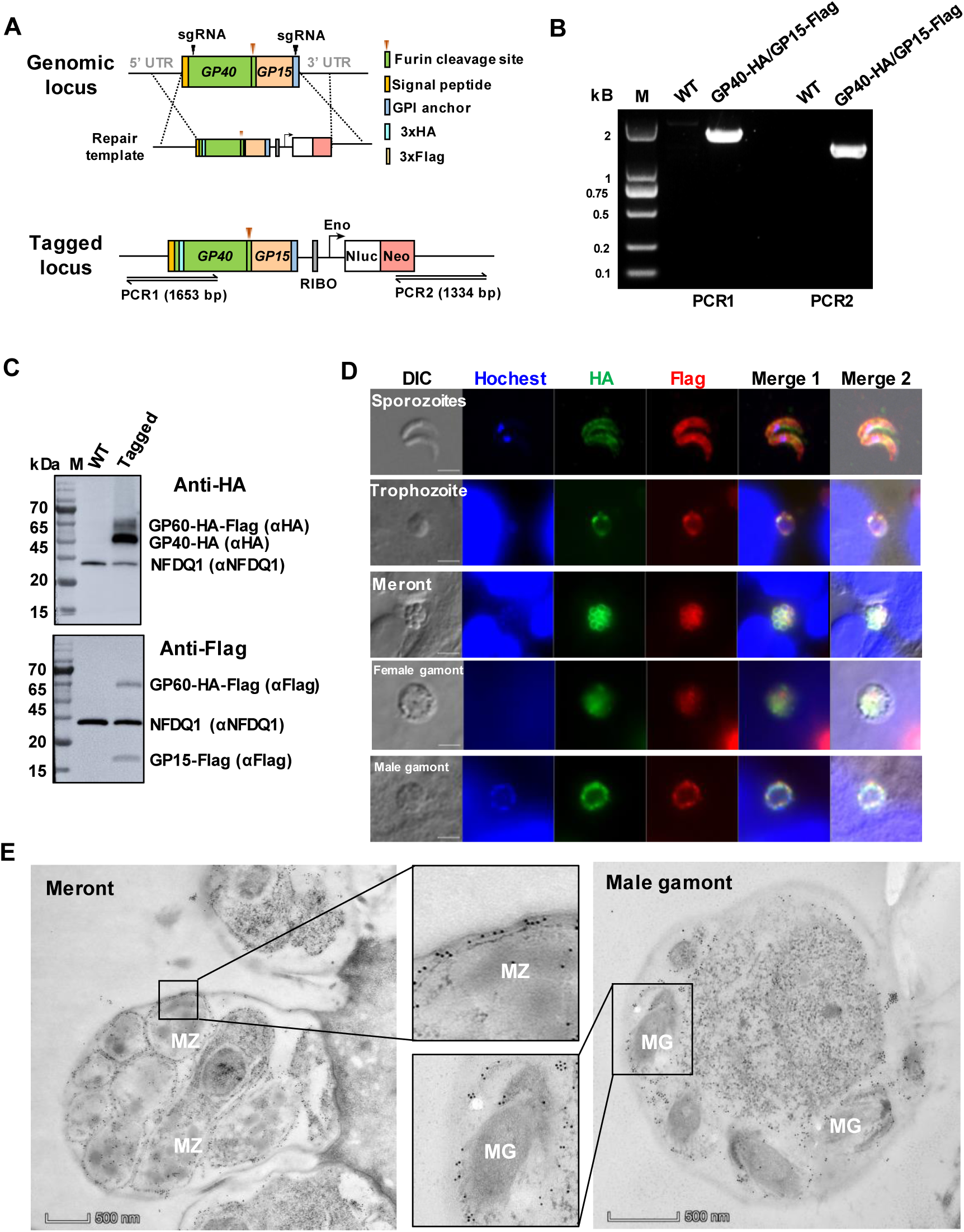
Endogenous gene tagging supports the membrane location GP60 protein expression in *Cryptosporidium parvum*. **A.** Strategy used in tagging the N-terminal end of the GP40 fragment with a 3× hemagglutinin (HA) epitope and the N-terminal end of the GP15 fragment with a 3× FLAG epitope using CRISPR/Cas9. **B.** Confirmation of the correct integration of the tagging cassette through PCR analysis of the GP40 fragment tagged with the HA sequence and the GP15 fragment tagged with the FLAG sequence. No product is generated in PCR analysis of the wildtype (WT). **C.** Confirmation of the correct tagging of the GP40 and GP15 products of GP60 by Western blot (WB) analysis of crude protein extracted from *C. parvum* oocysts using mAbs against the HA and FLAG tags using the expression of NFDQ1 (detected by a polyclonal antibody against recombinant NFDQ1) as control. As expected, both the GP60 precursor and the GP40 product are seen in WB analysis with the HA mAb, and both GP60 and the GP15 product are seen in WB analysis with the FALG mAb. No HA or FLAG-tagged products are seen in WB analysis of the WT. **D.** Expression of the GP40 (tagged with HA) and GP15 (tagged with FLAG) products of GP60 in free sporozoites and developing stages of *C. parvum* cultured in HCT-8 cells, as indicated by immunofluorescence analysis. GP40 and GP15 colocalize on the surface of sporozoites, later trophozoites, merozoites within mature meront, and gametes within male gamonts, but not on the surface of female gametes. **E.** Confirmation of the surface location of GP40 by immunoelectron microscope analysis of merozoites (MZ) within a meront and gametes (MG) within a male gamont using the HA mAb.

PCR analysis of the transfectant confirmed the correct integration of the tagging cassette into the genome (Fig. 3*B*). Results of Western blot analysis of oocyst lysate agreed with this finding; the mAb against HA reacted with two bands of ∼65 and 50 kDa, likely representing the HA-tagged GP60 and its furin cleavage product GP40, while the mAb against the FLAG tag detected the full GP60 protein and its furin cleavage product GP15 (Fig. 3*C*). IFA analysis of life cycle stages of the endogenously tagged mutant using mAbs against HA and FLAG showed abundant expression of GP40 and GP15 on surface sporozoites, trophozoites, merozoites and male gametes. In contrast, the female gametes showed low reactivity to both mAbs. In the IFA analysis, GP40 and GP15 colocalized with each other, allowing comparative studies of their fate during invasion and development of *C. parvum* (Fig. 3*D*). This result was confirmed by IEM analysis of the tagged line, which showed ample GP40 expression on the surface of these developmental stages (Fig. 3*E*, Fig. S1).

### GP60 is translocated to the feeder organelle during invasion and re-synthesized in early intracellular development

We examined the dynamics of GP60 expression using the endogenously tagged line and the mAb against the HA tag, which was incorporated into the N-terminal end of the GP60. During epithelial cell invasion, GP60 (or its GP40 product) was translocated from the sporozoite surface to the apical end (Fig. 4*A*). In enveloped sporozoites and newly formed trophozoites, GP60 expression was largely restricted to the feeder organelle, with no visible detection of the protein in the remaining parts of the developing parasites by IEM analysis (Fig. 4*B*).

**Figure 4.**
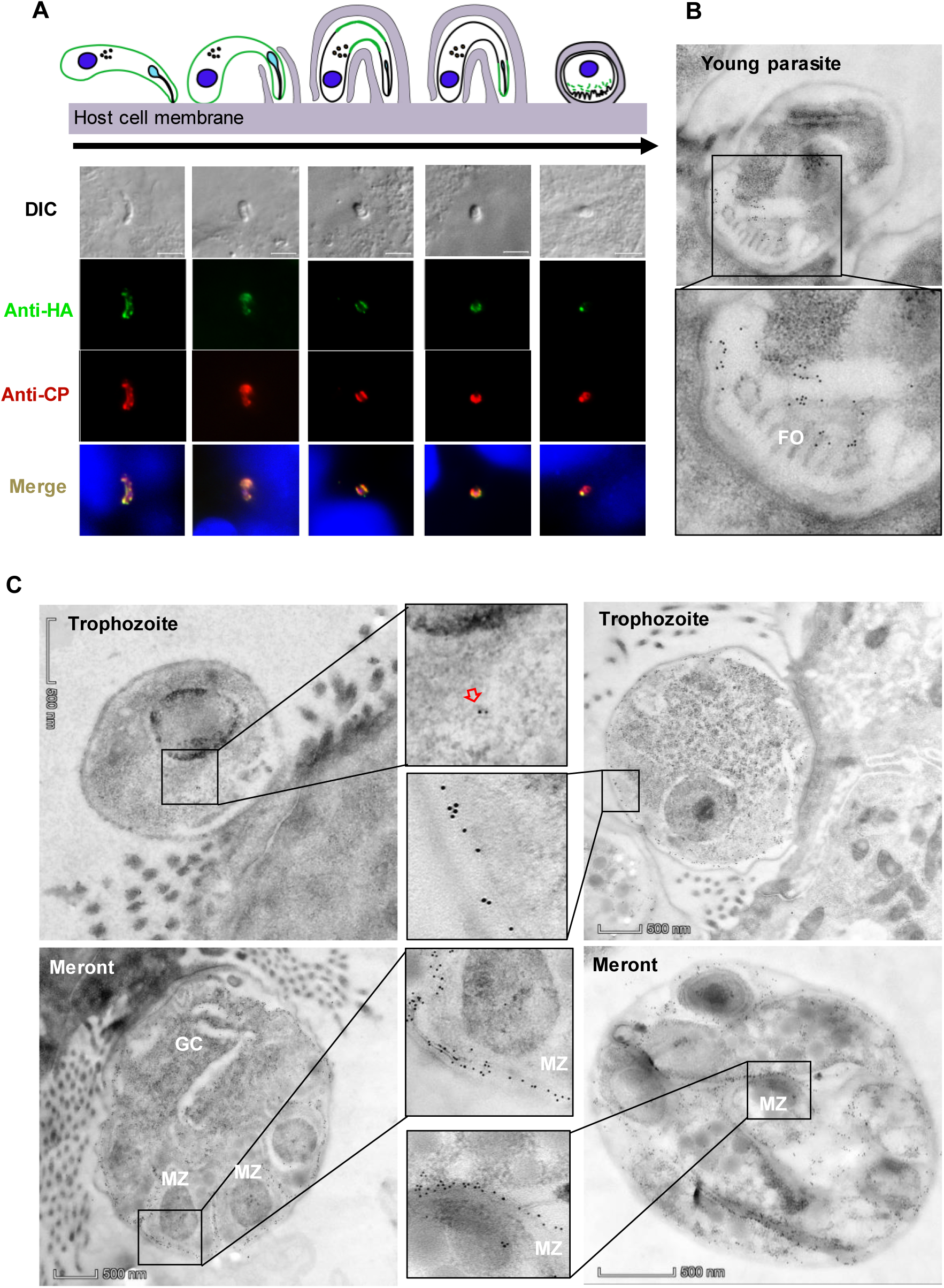
GP60 is shed during invasion and resynthesized during development. **A.** Movement of GP60 during invasion of host cells by *Cryptosporidium parvum* sporozoites as indicated by the HA-tagging of the GP40 product of GP60, using a mAb against the HA tag to track GP40 and a polyclonal antibody against sporozoites (Sporo-Glo) as a control (Anti-CP). GP40 moves to the apical end of the sporozoite and concentrates on one side as the sporozoite enters the host cell and rounds up. **B.** Confirmation of the localization of GP40 at the *C. parvum*-host interface (the feeder organelle, FO) by immunoelectron microscopy using the HA mAb. **C.** GP60 is resynthesized in trophozoites and moves to the surface of later trophozoites and merozoites (MZ) within mature meronts. During the early *C. parvum* development in host cells, only a few gold particles are present in the cytosol of newly formed trophozoites (red arrow). Gold particles increase in number and are mostly seen on the surface of late trophozoites. In 4N meronts, gold particles are mostly seen on the surface of newly formed merozoites (MZ) and within the germinal center (GC). Once the 4N meronts mature to 8N, gold particles are seen only on the surface of the merozoites.

RNA-seq analysis indicated that the *GP60* gene was not expressed in oocysts and newly excysted sporozoites. Soon after invasion, *GP60* gene expression was first detected, gradually peaking at 12 h and remaining at this level at 24, 36, 48 and 72 h (Fig. 1*D*). Consistent with this timing, IEM examinations of HA-tagged mutant indicated that GP60 was initially expressed in the cytosol of young trophozoites and subsequently translocated to the surface of mature trophozoites. In 4N meronts, GP60 was mainly found on the surface of developing merozoites and in the germinal center where new merozoites budded out. As meronts mature to 8N, GP60 was present only on the surface of merozoites (Fig. 4*C*).

### GP60 functions as both a secretory and a membrane protein

Since GP60 has both a signal peptide and a GPI anchor, we examined the effect of each domain on the translocation of the protein (Fig. 5*A*). The successful construction of these mutant lines was confirmed by PCR analysis using primers franking the insert and 5′and 3′sequences of the native locus, producing the expected PCR products in mutant lines only (Fig. S2*A* and Fig. S2*B*). Depletion of the signal peptide prevented the protein from being secreted to the parasite surface. As a result, the synthesized GP60 accumulated in the cytosol of sporozoites, trophozoites and merozoites, as shown by IFA analysis using mAbs against the incorporated HA and FLAG tags (Fig. 5*B* and 5*C*). In contrast, depletion of the GPI anchor sequence resulted in the segregation of newly synthesized GP60 into the PV space between parasite and host membranes (for trophozoites, meronts and male gamonts) and into the space between merozoites (for mature meronts) (Fig. 5*B* and 5*D*). This result was confirmed by comparative IEM analysis of the tagged wildtype (WT) and mutant lines (Fig. S1).

**Figure 5.**
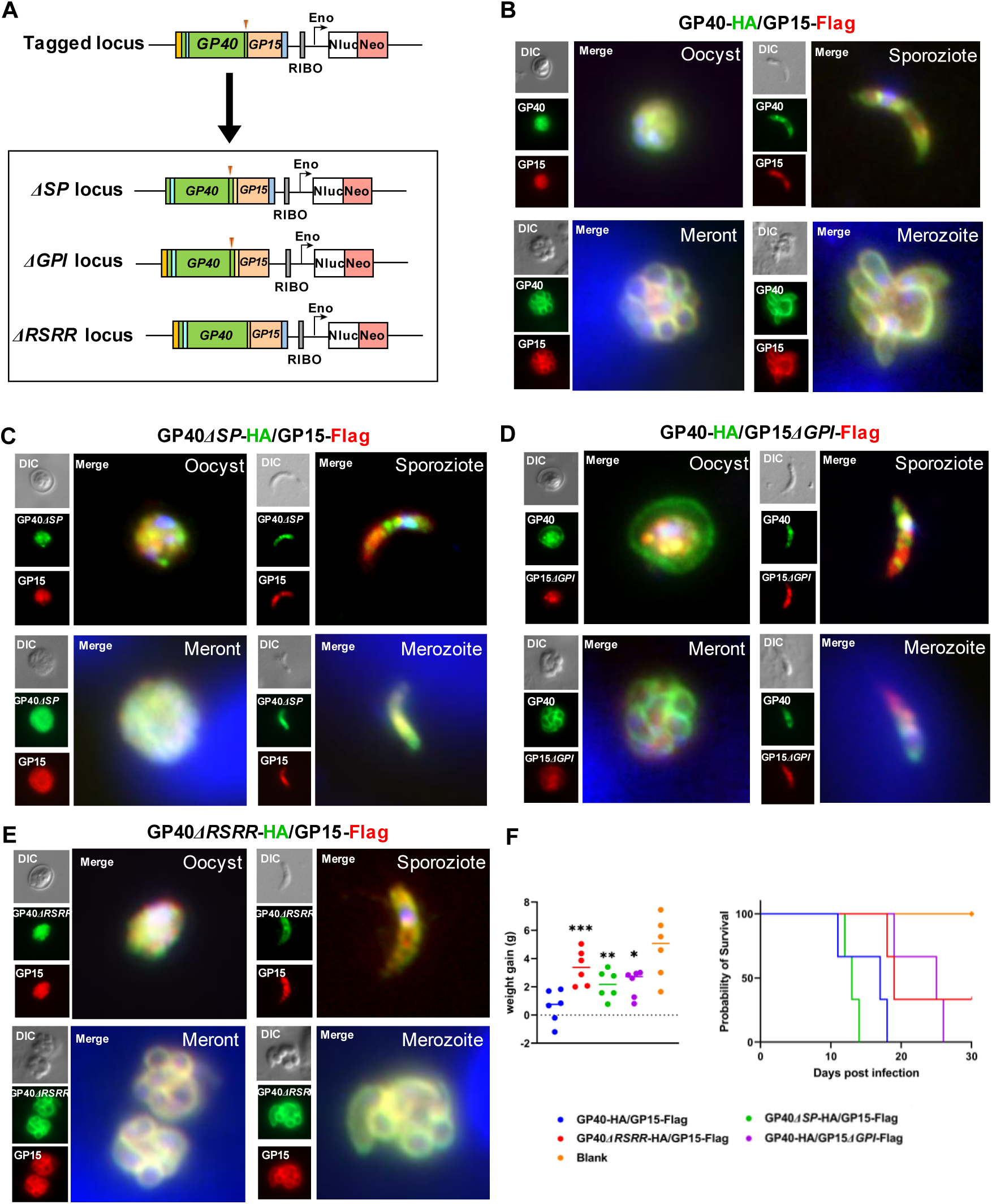
GP60 functions as both a secretory and a membrane protein. **A.** Strategy used in the depletion of the signal peptide (SP), glycophosphatidylinositol (GPI) anchor, and furin cleavage (RSRR) sequences in *Cryptosporidium parvum* using CRISPR/Cas9. The GP40-HA/GP15-FLAG construct in Fig. 2A is used as the template in further modifications of the *GP60* gene. **B.** Immunofluorescence analysis (IFA) of the expression of the GP40 and GP15 products of GP60 in transgenic *C. parvum* that has been endogenously tagged at the *GP60* locus as shown in Fig. 2A, using mAbs against HA and FLAG. As expected, GP40 (tagged with HA) and GP15 (tagged with FLAG) are expressed on the surface of sporozoites and merozoite and colocalize with each other. **C.** IFA of the expression of GP40 and GP15 products of GP60 in transgenic *C. parvum* with the signal peptide depleted from GP60. GP40 and GP15 are now expressed in the cytosol and do not colocalize with each other. **D.** IFA analysis of the expression of GP40 and GP15 products of GP60 in transgenic *C. parvum* with the GPI anchor depleted from GP60. GP40 and GP15 loss the membrane expression and do not colocalize with each other. In particular, GP40 is also detected on the oocysts wall and accumulates in the space between merozoites within meronts. **E.** IFA analysis of the expression of GP40 and GP15 products of GP60 in transgenic *C. parvum* with the furin cleavage site depleted from GP60. GP40 and GP15 expression is mostly similar to that in Fig 5a. **F.** Body weight (DPI 2-DPI 12) and survival (DPI 30) of IFN-γ knockout mice infected with mutant *C. parvum* lines described above as determined at the end of the infection study (n = 3).

After depletion of the signal peptide or GPI anchor, GP40 (tagged by HA) and GP15 (tagged by FLAG) were no longer co-localized, especially during the oocyst stage (Fig. 5*C* and 5*D*). In oocysts, GP60 was normally expressed only on sporozoites, with colocalization of the GP40 and GP15 products of the processed protein. However, in oocysts of mutants with a depleted GPI anchor in the *GP60* gene, GP40 was expressed not only inside sporozoites but also in the oocyst space, while GP15 was restricted to inside sporozoites (Fig. 5*D*). This was particularly evident in the IEM analysis of the mutants with the HA mAb, which showed the presence of gold particles in oocysts space and the space between oocyst wall and PVM (Fig. S1). These data suggested that GP40 was secreted outside the mature stages and anchored to the parasite surface by GP15.

### GP60 is proteolytically processed

We replaced the native *GP60* gene of *C. parvum* with a *GP60* sequence that had no furin cleavage site (with the RSRR sequence). Targeted sequencing confirmed successful removal of the 12 nucleotides encoding the cleavage sequence (Fig. S3*A*). Western blot analysis of crude sporozoite proteins using mAbs against the HA and FLAG tags indicated that the depletion of the RSRR sequences prevented cleavage of GP60 into GP40 and GP15 (Fig. S3*B*). However, this lack of processing did not affect the translocation of the GP60 protein, since both GP40 (tagged with HA) and GP15 (tagged with FLAG) co-localized on the surface of sporozoites and merozoites in IFA analysis (Fig. 5*E*). This result was also confirmed by IEM analysis with the HA mAb (Fig. S3*C*).

### Disrupted translocation and processing of GP60 affect the growth and host infectivity of *C. parvum*

Depletion of the signal peptide, GPI anchor, or furin cleavage sequences had some effect on the growth and pathogenicity of *C. parvum*. During *in vitro* infection of HCT-8 cells, the depletion of the signal peptide reduced the growth of *C. parvum* at 24, 36 and 48 h, although no obvious effect was observed when the GPI anchor and furin cleavage sequences were deleted (Fig. S4*A*). However, the depletion of the GPI anchor and furin cleavage sequences moderately reduced the parasite load in infected GKO mice as indicated by fecal luciferase measurements and oocysts per gram of feces (OPG, Fig. S4*B*). In addition, mice infected with these mutants had a higher body weight than those infected with the HA-tagged WT line (Fig. 5*F*), and those infected with mutants depleted of the GPI anchor or furin cleavage sequences survived longer (Fig. S4*B*).

Histopathological examination of the ileum was consistent with this observation; mice infected with these mutants had lower parasite load, fewer changes in intestinal architecture, and a higher villus height to crypt depth ratio than those infected with the HA-tagged WT line (Fig. S4*C*, Fig. S4*D*).

### GP60 is dispensable

To assess the biological significance of GP60, we replaced the *GP60* gene sequence in the *C. parvum* genome with the mCherry sequence (Fig. 6*A*). Surprisingly, GP60 was dispensable as we were able to generate a mutant expressing mCherry but lacking the *GP60* sequence of (Fig. 6*B*). The complete deletion of the gene was confirmed by PCR analysis of the mutant (Fig. 6*C*). The oocyst shedding of the mutant was initially low, but increased significantly after a secondary passage in mice (Fig. 6*D*). Immunofluorescence analysis of intracellular stages at different cultivation time with the GP40 mAb confirmed complete GP60 depletion (Fig. 6*E*). WGS analysis of the mutant showed no duplication of the *GP60* gene elsewhere in the genome. The gene depletion had no visible effect on *C. parvum* morphology, as developmental stages of the GP60- KO and GP60-tagging (GP40-HA/GP15-Flag) lines looked similar in SEM analysis of developmental stages in HCT-8 cells (Fig. 5).

**Figure 6.**
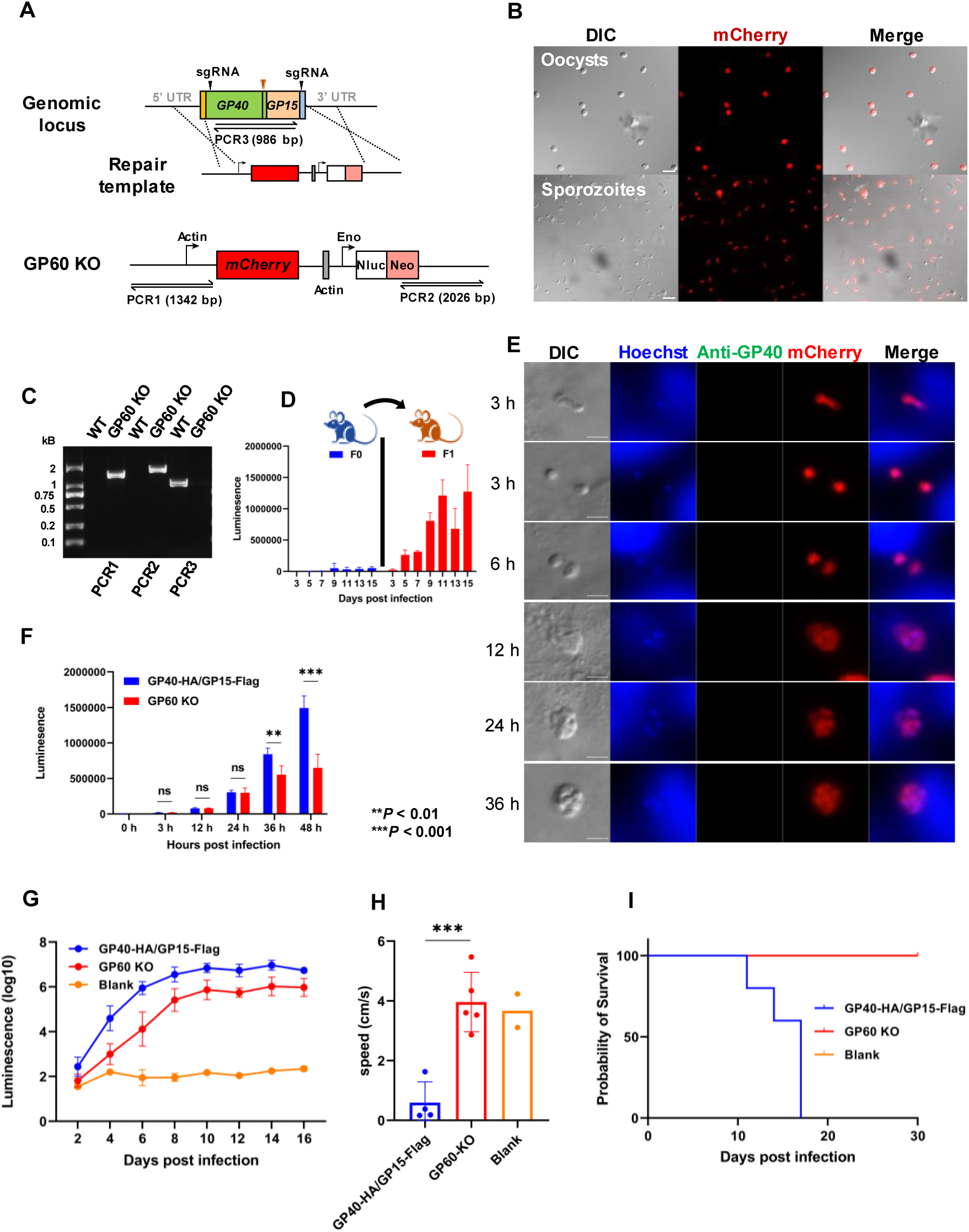
GP60 in *Cryptosporidium parvum* is dispensable. **A.** Strategy used in the replacement of the *GP60* sequence with sequences encoding mCheery, NanoLuciferase (Nluc), and neomycin resistance gene (Neo). **B.** Microscopy of oocysts from mice infected with the transgenic *C. parvum*, with almost all oocysts being red. **C.** Confirmation of the correct integration of the knockout cassette through PCR analysis. No product is generated in PCR analysis of the wildtype (WT), except for PCR 3 conducted using *GP60*-specific primers. **D.** Serial passages of the transgenic line in mice to increase the infection intensity for the characterization of the transgenic line. **E.** Confirmation of the depletion of GP60 through immunofluorescence analysis using a mAb against GP40. No GP60 is expressed on mCherry-positive parasites in HCT-8 cultures. **F.** Effect of GP60 depletion on the growth of *C. parvum* in HCT-8 cells in comparison with the HA and FLAG-tagged WT (control) line. **G.** Differences in the infection intensity of *C. parvum* as measured by fecal luciferase activity between IFN-γ knockout (GKO) mice infected with the GP60-depleted line and those infected with the HA and FLAG-tagged WT line. **H.** Comparison of physical activity between GKO mice infected with GP60-depleted and control lines. **I.** Differences in the survival of GKO mice infected with GP60-depleted and control lines.

### GP60 plays important role in host infectivity of *C. parvum*

The GP60 depletion had a profound effect on the growth of *C. parvum*. The GP60-KO mutant grew slower than the HA-tagged WT line, especially during late development *in vitro* (Fig. 6*F*). In GKO mice, the mutant line produced a lower infection intensity than the HA-tagged line, as indicated by the measurement of fecal luciferase activity and OPG (Fig. 6*G* and Fig. S6*A*). As a result, mice infected with the GP60-KO mutant gained weight faster than those infected with the WT line. Moreover, they showed no overt clinical signs of infection, and exhibited normal movement and fecal output. In contrast, mice infected with the HA-tagged line showed lethargy, significant weight loss, and reduced fecal output due to inappetence (Fig. 6*H*, Fig. S6*B*, Fig. S6*C*, Fig. S7). Consistent with this, mice sacrificed at the peak of *C. parvum* infection showed differences in parasite load between the two groups. Notably, mice infected with the GP60-KO mutant showed lower numbers of parasites on the surface of the ileum than those infected with the HA-tagged line (Fig. S4*D* and S4*E*). They maintained a largely normal villus structure, with a significantly higher villus height/crypt depth ratio than those infected with the HA-tagged line (Fig. S6*F*). All mice infected with the GP60-KO mutant survived the infection, while those infected with the HA-tagged lined succumbed by DPI 18 (Fig. 6*I*).

### GP40 and GP15 are not individually critical for *C. parvum* infection

Mutants in which the GP40 or GP15 fragment of the *GP60* gene were removed were also successfully generated *in vivo* (Fig. S8*A* and 8*B*). The correct integration of the gene deletion cassettes was confirmed by allele-specific PCR using primers franking the native and insert sequences (Fig. S2*C* and S2*D*). This was further confirmed by microscopy, as oocysts with GP40 or GP15 depleted were red due to the replacement of gene fragments encoding them with the mCherry sequence. IFA analysis of developmental stages using the mAb against GP60 and a polyclonal antibody against GP15 further confirmed the loss of GP40 or GP15 expression in these mutants (Fig. S8*C*).

The gene deletion changed the location of the two GP60 fragments. Although oocysts generated by the GP40-KO and GP15-KO mutants were red due to the replacement of the fragments with mCherry, sporozoites excysted from them were colorless, indicating that the association between GP40 and GP15 on the sporozoite membrane was prevented by the replacement of the GP40 or GP15 with mCherry (Fig. S8*B*). This was expected, as the GP40-KO and GP15-KO mutants both had kept the furin cleavage site, which would hydrolyze the protein expressed by the GP40-KO and GP15-KO mutants to mCherry and GP40 or GP15. Similarly, the GP40 depletion changed the location of GP15 expression in mature meronts from the membrane to the cytosol, probably due to the lack of signal peptide after the furin cleavage. In addition, the GP15 depletion changed the location of GP40 in mature meronts from membrane of merozoites to space between them due to the loss of the GPI anchor after the furin cleavage (Fig. S8*C*, Fig. S9*A*, Fig. S9*B*). As a result, GP40 and the mCherry did not co-localize with each other, with mCherry remaining inside merozoites (Fig. S9*A*, Fig. S9*C*).

The depletion of GP40 and GP15 products individually had no apparent effect on *C. parvum* growth *in vitro*. In addition, GKO mice infected with the GP40 or GP15 depleted mutant and the GP60-tagging line had similar oocyst shedding the throughout the course of *C. parvum* infection. Consistent with this, the depletion of GP40 or GP15 had no effect on body weight gain and survival of infected GKO mice. In contrast, as observed above, mice infected with the GP60-KO mutant had significantly lower oocyst shedding, higher bodyweight and longer survival than those infected with the GP60-KO line (Fig. S8*E*-*H*). However, in the less susceptible 4-week-old WT C57BL/6 mice, the depletion of GP40 or GP15 reduced *C. parvum* infection intensity as measured by luciferase activity and OPG during the early infection period (DPI 4-DPI 8), although it had no effect on the bodyweight of infected mice (Fig. S10).

### GP60 contributes to host infectivity

Previously, we showed that the IIdA20G1-HLJ isolate induced significantly higher oocyst shedding intensity and duration than the IIaA17G2R1-IOWA isolate, leading to lower body weight gain and survival of infected GKO mice (20). To further evaluate the role of GP60 in host infectivity of *C. parvum*, we replaced the *GP60* gene in the genome of the highly infective IIdA20G1-HLJ isolate with the *GP60* sequence of the avirulent IIaA17G2R1-IOWA isolate (Fig. 7*A*). The gene replacement was confirmed by the ectopic expression of mCherry in transgenic oocysts as revealed by fluorescence microscopy (Fig 7*B*), and by DNA sequence analysis (Fig. S2*E*). It resulted in slower *in vitro* growth of *C. parvum* during the sexual phase of development (Fig. 7*C*). It also reduced the intensity of oocyst shedding in GKO mice compared to those infected with the tagged WT line (Figure 7*D*). In addition, mice infected with the replaced *GP60* gene survived longer (Fig. 7*G*). Consistent with this, mice infected with the IIa GP60 mutant had fewer parasites on the ileal surface and less damage to the villus architecture than those infected with the tagged line carrying the WT IId *GP60* gene (Fig. 7*E* and 7*F*).

**Figure 7.**
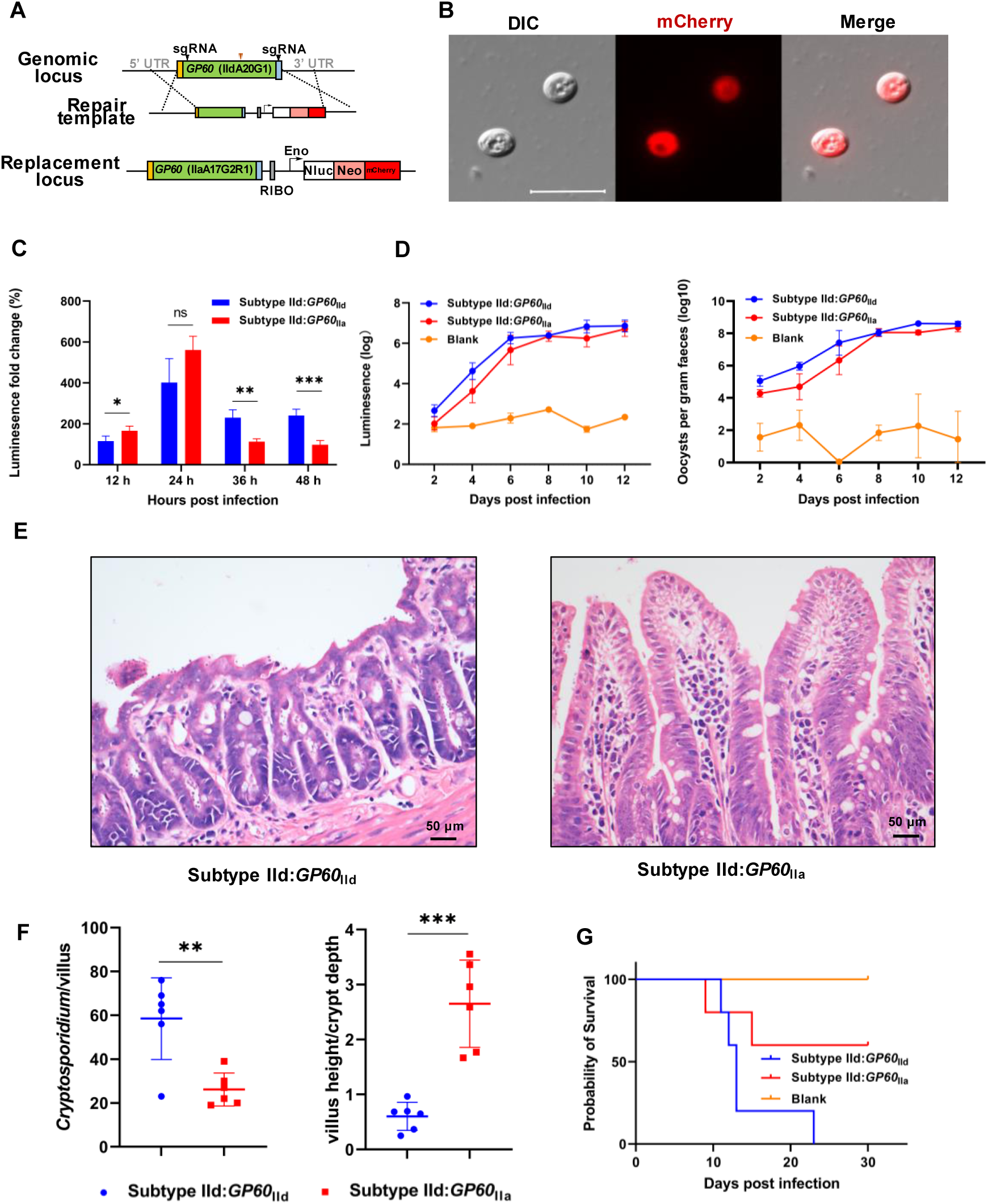
GP60 contributes to host infectivity as indicated by gene replacement results. **A.** Strategy used in the replacement of the *GP60* gene in the virulent IIdA20G1-HLJ isolate with the *GP60* sequence from the a virulent IIaA17G2R1-IOWA isolate, with the incorporation of sequences encoding mCheery, NanoLuc luciferase (Nluc), and neomycin resistance gene (Neo) for the identification and selection of transgenic parasites. **B.** Microscopy of oocysts from mice infected with the transgenic line, with the transgenic oocysts being red. **C.** Effect of the GP60 replacement on the growth of *C. parvum* in HCT-8 cells in comparison with those infected with the HA and FLAG- tagged WT line. The gene replacement has reduced the growth of *C. parvum* at 36 h and 48 h. **D.** Effect of the GP60 replacement on the infection intensity of *C. parvum* in IFN-γ knockout (GKO) mice as measured by fecal luciferase activity and oocysts per gram of feces, in comparison with the uninfected control (the same group of mice used in Fig. S8). **E. & F.** Differences in the ileum architecture (*E*) and parasite load and villus height to crypt depth ratio (*F*) in the ileum between mice infected with the virulent WT line (Subtype IId) and those infected with the GP60 replacement line (Subtype IId:GP60IIa). Mice infected with the latter have fewer parasites and less intensive damage in the ileum. **G.** Differences in the survival of GKO mice after infection with the GP60 replacement line (Subtype IId:GP60IIa) and similarly tagged WT line (Subtype IId).

## Discussion

One of the long-standing issues in cryptosporidiosis research is the parasite-host interaction during early infection (21). More than a handful of parasite molecules located on the apical end and surface of sporozoites have been implicated in the initial attachment and invasion process of *C. parvum* (22). They include several members of the mucin glycoprotein family such as Muc4, Muc5, GP900, and GP60. Of these, GP60 and its cleavage products GP40 and GP15 have received the most attention. Studies using neutralizing mAbs have implicated GP60 and its GP40 and GP15 products in sporozoite and merozoite attachment to and invasion of enterocytes (8, 23). Differences in host preference and virulence have been reported between GP60 subtype families of *C. parvum* and *C. hominis*, respectively (14, 15). Mice experimentally infected with different GP60 subtypes of *C. parvum* have shown significant differences in infection intensity, duration, and pathogenicity (20). However, few studies have been undertaken to examine the biological processes of GP60-mediated host cell infection. Underlying the poor understanding of *Cryptosporidium* infection is the lack of in-depth studies of the parasite-host interactions using advanced genetic tools (22). CRISPR/Cas9-based genetic manipulation tools have only recently been used by one research group to systematically characterize the role and process of ROP and MEDLE proteins in early *C. parvum*-host interactions (24, 25). These proteins have been shown to be exported into host cells and to modify host cytoskeletal and other responses during *C. parvum* invasion and infection process.

Data from this study suggest that the GP60 protein likely plays multiple roles in *C. parvum*-host interactions and contributes to host infectivity. Consistent with its immunodominant nature, GP60 is expressed on the surface of mobile stages (sporozoites, merozoites, and male gametes), suggesting that GP60 may be involved in sporozoite and merozoite invasion and fertilization of *C. parvum*. Once synthesized in the parasite cytosol, GP60 is secreted outside the developing stages through its signal peptide, cleaved at its furin cleavage site into GP40 and GP15, anchored to the surface of *C. parvum* developing stages through the GPI anchor at the C-terminal end of GP15, translocated to the apical end of the sporozoites during invasion, and remains detectable only at the pathogen-host interface. While the GP40 and GP15 products of GP60 depend on each other for translocation to the parasite surface, the full GP60 protein is required for its involvement in host infectivity.

We have confirmed the surface location of the GP60 protein. Analyses using mAb staining and endogenous gene tagging have both shown high levels of GP60 on the surface of sporozoites, merozoites, and male gametes. Previously, several studies using mAbs have shown the expression of GP60 on the surface of both sporozoites and merozoites (6, 8, 17). We believe that the previous suggestion of GP60 as a PVM protein (18) was the result of a misinterpretation of the surface location of GP60 in late trophozoites and merozoites under immunofluorescence microscopy. We have furthered our understanding of GP60 expression by demonstrating that the *GP60* gene is the most highly expressed gene in the *C. parvum* genome by transcriptome analysis of both *in vitro* and *in vivo* infections. Further studies are needed to determine whether GPI-anchored GP60 is transported from the Golgi to the plasma membrane via vesicles following the conventional sorting system for GPI-anchored proteins (26), or is first transported into micronemes where GP60 has been identified (19).

More importantly, we have shown for the first time that GP60 is highly expressed on the surface of male gametes. To date, only a few proteins, such as α-tubulin and HAP2, are known to be expressed at high levels in male gametes of *Cryptosporidium* spp. (27). Like α-tubulin, GP60 is also expressed in sporozoites and merozoites, but rarely in female gametes. The function of GP60 in male gametes is not yet known, but the fact that GP60 is involved in the initial parasite-host interaction and the surface location of the protein suggest that the protein may be involved in the fertilization of female gametes by the male gametes. This is further supported by the observation that the suppression of parasite growth *in vitro* by GP60 gene depletion and replacement mainly occurred during the sexual development of *C. parvum*. Currently, little is known about the sexual development in *Cryptosporidium* spp. As blockage of male gamete entry into female gametes has been suggested as a cause of incomplete development of *C. parvum in vitro*, sexual reproduction likely plays an important role in pathogen transmission (27). The identification of a surface protein in male gametes opens a new avenue for studies of sexual reproduction in *Cryptosporidium* spp. To date, studies on gene expression in sexual stages have been limited to female gametes due to lack of surface markers for flow cytometric sorting of male gametes (19, 27).

In agreement with the multifunctional nature of GP60 protein, the expression of the *GP60* gene undergoes dynamic changes during the life cycle of *C. parvum*. While the GP60 protein is highly expressed in sporozoites, the gene encoding it is not transcribed during this stage. During invasion, GP60 moves to the apical end and is shed by sporozoites and merozoites gliding on host cells (28). Shortly after the entry into the host membrane, GP60 is detectable only at the parasite-host cell interface, reinforcing its role during the invasion. Once the newly invaded parasite is rounded off, GP60 is no longer detected, and new GP60 is synthesized in the cytosol of developing trophozoites and translocated to the surface of late trophozoites in preparation for merozoite biogenesis. Both the signal peptide and GPI anchor sequences are required for GP60 translocation, as depletion of the signal peptide resulted in GP60 accumulation in the cytosol, and depletion of the GPI anchor sequence changed the membrane location of the GP40 product to free in the parasitophorous vacuole and oocyst. This finding is expected because in eukaryotic cells the signal peptide is required for proper sorting of secretory proteins in the endoplasmic reticulum (29). Similarly, the GPI anchor is required for the sorting of GPI-anchored proteins from other secretory proteins in the trans-Golgi network for transport to the plasma membrane (26).

The surface location of GP60 is consistent with its highly variable and immunodominant nature. Since its initial discovery, the *GP60* gene has been known as the most polymorphic gene in the *Cryptosporidium* genome, expressing different families of GP60 proteins with highly divergent sequences (6, 30). *C. parvum* isolates from different subtype families differ from each other in their host preference, with IIa being mainly found in calves and humans, IId mainly in lambs, goat kids, and humans, and IIc almost exclusively in humans (14). In contrast, *C. hominis* families appear to differ from each other in virulence, with the Ib subtype family, particularly the IbA10G2 subtype, causing more severe clinical manifestations than other subtype families in some epidemiological studies (15). Therefore, GP60 characteristics have been linked to the phenotypic characteristics of *Cryptosporidium* isolates. This correlation is also supported by the dominance of the *C. parvum* IIaA15G2R1 subtype worldwide and the *C. hominis* IbA10G2 subtype in Europe and other high-income countries (31, 32). As one of the two most dominant antigens in *C. parvum* and *C. hominis* (33, 34) and one of the few protective *Cryptosporidium* antigens against reinfection (12), immune and other adaptive selection has likely occurred in this gene (35, 36).

In this study, we provide experimental evidence for GP60 as a key factor in *C. parvum* infectivity and pathogenicity. Despite the biological importance of GP60, the gene encoding it is dispensable, suggesting the existence of alternative mechanisms for zoite invasion. However, depletion of the *GP60* gene significantly reduced the growth of *C. parvum*, and GKO mice infected with the mutant line showed significantly lower parasite loads and no clinical signs of infection, while mice infected with the tagged WT line experienced severe infection and exhibited lethargy, arched backs, wet feces, and weight loss. All mice infected with the WT line died, while those infected with the mutant line survived the infection. More importantly, replacing the IId *GP60* sequence with the IIa sequence from an avirulent isolate significantly attenuated the WT line, supporting the involvement of GP60 in host infectivity.

Surprisingly, although the GP40 product of GP60 is widely thought to be involved in zoite attachment to host cells and relies on the GP15 product for membrane localization, the full GP60 protein appears to be required for GP60-mediated host infectivity. Depletion of the *GP40* or *GP15* fragments of the *GP60* gene alone has only a minor effect on *C. parvum* growth and pathogenicity. Similarly, disruption of the translocation and processing of the GP60 proteins by deleting the signal peptide, GPI anchor and furin cleavage sequences only moderately reduces the infectivity and pathogenicity of the pathogen. This, together with the disposable nature of the complete *GP60* gene, suggests that there are compensatory mechanisms for GP60-mediated infection of host cells. Although yet observed in *Cryptosporidium* spp., plasticity and functional redundancy are common features of gene families involved in the invasion process of *Plasmodium* spp. and *T. gondii* (37, 38).

In conclusion, we have performed the first biological study of a major immunodominant *Cryptosporidium* antigen using reverse genetic tools. The data obtained suggest that GP60 likely plays multiple functions during *C. parvum* invasion and reproduction and is an important factor in host infectivity. Further studies are needed to identify the host cell receptors involved and the mechanisms underlying GP60-mediated functions. Such studies should also evaluate the potential for the use of the GP60-depleted line as a vaccine candidate against *C. parvum*.

## Materials and Methods

### Ethics statement

The laboratory animals used in this study were housed and treated in accordance with the Regulations for the Administration of Affairs Concerning Experimental Animals of the People’s Republic of China. All experiments were conducted under the approval of the Laboratory Animal Research Center of Guangdong Province and South China Agricultural University (approval number: 2021C005).

### Mice

IFN-γ knockout (GKO)and wild-type C57BL/6J mice were purchased from Jackson Laboratories (Bar Harbor, Maine, USA) and bred at the Laboratory Animal Center of South China Agricultural University. Mice were housed in individually ventilated cages and provided with sterilized food, water, and bedding. Most of the infection studies were performed on mice aged 3-5 weeks.

### HCT-8 cell culture and infection

The human ileocecal adenocarcinoma cell line HCT-8 (ATCC CCL-244) was obtained from the Chinese Academy of Sciences for *in vitro* cultivation of *C. parvum*. HCT-8 cells were seeded in 24-well plates and grown to approximately 80% confluence. The cells were then inoculated with*C. parvum* oocysts in RPMI 1640 culture medium (Gibco, Grand Island, USA) supplemented with 2% FBS (Gibco, Grand Island, USA) as described (Xu et al., 2019). Invasion stages of parasites were obtained by infecting HCT-8 cells for 5 to 30 min, while intracellular stages of parasites were obtained by infecting HCT-8 cells for 3 to 48 h. Monolayers were harvested at different time to observe trophozoites (3 h), meronts (12, 24, and 24 h), and male and female gametes (48 h) using the IFA assay.

### Oocyst preparation and excystation

The *C. parvum* IIdA20G1-HLJ isolate was obtained from dairy calves in Heilongjiang (HLJ) Province, China (20), while the *C. parvum* IIaA17G2R1-IOWA isolate was purchased from Waterborne, Inc. (New Orleans, USA). They were passaged in calves or GKO mice, with oocysts being purified as described (20), and stored at 4°C in PBS for up to 6 months before use. Prior to infection, oocysts were treated with 10% Clorox Household Bleach (Clorox, Oakland, USA) on ice for 10 min, followed by centrifugation at 13,200 rpm and 4°C for 3 min. After discarding the supernatant, the oocysts were washed three times with PBS. The oocysts were then resuspended in 400 μL of 1% BSA-PBS and mixed with 400 μL of 0.2 mM sodium taurodeoxycholate solution (Sigma-Aldrich, Saint Louis, USA) to a final concentration of 0.75%. The excystation mixture was incubated for 1 h at 37°C in a water bath, and the resulting sporozoites obtained were washed three times by centrifugation at room temperature with PBS.

### Transcriptome analysis

For *in vivo* experiments, GKO mice were infected with 10,000 *C. parvum* oocysts and their intestines were harvested at DPI 12 for RNA extraction. For *in vitro* experiments, HCT-8 cells were seeded in 48-well plates and cultured as described above, followed by infection with 4×10^5^ bleached *C. parvum* oocysts per well. RNA extraction from free sporozoites was performed immediately after oocyst shedding, while RNA extraction from parasite-infected cells was performed at 3, 6, 12, 24, 36, 48 and 72 h post infection. Four replicates were performed for each time period and negative controls consisted of uninfected cells. The extracted RNA was sent to Genedenovo Biotechnology Co., Ltd (Guangzhou, China) for transcriptome sequencing using the Illumina technology. FastQC (v0.11.9, https://github.com/s-andrews/FastQC) was used for quality control of the sequence reads, while STAR (2.7.9a) was used to map the raw reads to the *C. parvum* IIdA20G1-HLJ reference genome (SRA, SRR15694560). FPKM values for the gp60 and reference genes were calculated using RSEM (v1.3.1, https://github.com/deweylab/RSEM.git), and gene expression levels were visualized as violin plots using the R package ggplot2 and Manhattan plots using the R package qqman. The RNA-seq data generated in the study have been submitted to NCBI under BioProject No. PRJNA1011005.

### Extended methods

Other methods and procedures used in this study, including genetic manipulation of parasites using CRISPR/Cas9, Western blot analysis of proteins, characterization of parasites by immunofluorescence and electron microscopy, measurement of parasite load by luciferase assays and qPCR, and measurement of physical activity of infected mice, are described in detail in the online supplementary materials.

### Statistics

Statistical analyses were performed using GraphPad Prism 9.0 (GraphPad software). An unpaired t-test was used to compare the means of two groups at the same time point. Differences with *P* values <0.05 were considered significant.

## Acknowledgments

This work was supported in part by National Natural Science Foundation of China (32030109, 31972697, 31820103014, and 32150710530), the Guangdong Major Project of Basic and Applied Basic Research (2020B0301030007), 111 Project (D20008), and Double First-class Discipline Promotion Project (2023B10564003). We thank Jilei Huang, Chuanhe Liu and Xiaoxian Wu from the Instrumental Analysis & Research Center, South China Agricultural University for technical assistance.

