## Supplemental materials for "Variant Surface Protein GP60 Contributes to Host Infectivity of *Cryptosporidium parvum*"

### Supporting Information Text

#### Material and Methods

#### Histological examination

Intestines were harvested from parasite-infected GKO mice. Fragments of the intestines were fixed in 4% paraformaldehyde for 12 h, embedded in paraffin, and sectioned. The specimens were processed for H&E staining with hematoxylin and eosin, immunofluorescence staining with anti-GP40 primary antibody and Alexa Fluor-conjugated secondary antibodies, and immunohistochemistry with anti-GP40. The stained slides were examined under an Olympus BX53 microscope (Olympus, Tokyo, Japan).

#### Luciferase assay.

Luciferase assays were performed using the Nano-Glo Luciferase assay kit (Promega, Madison, USA) as previously described (1). For mouse fecal samples, one fecal pellet was collected in a 1.5 mL microcentrifuge tube to which eight 3-mm glass beads (Fisher Scientific, Hanover, USA) and 0.8 mL fecal lysis buffer (50 mM Tris pH 7.6, 2 mM DTT, 2 mM EDTA pH 8.0, 10% glycerol, 1% Triton X-100 prepared in water) were added. The tube was vortexed for 1 min and centrifuged at 19,000g for 1 min. Subsequently, 50 μL supernatant was added to a 96-well plate, followed by the addition of 50 μL of a 25:1 Nano-Glo Luciferase Buffer to Nano-Glo Luciferase Substrate Mix. The plate was then incubated for 3 min at room temperature in the dark.
For luciferase analysis of HCT-8 cell cultures, the medium was replaced with 100 μL Nano-Glo Luciferase Buffer, and the mixture was incubated at 37°C for 10 min. Afterwards, the cells were scraped, the resulting lysate was transferred to 96-well plates, and 1 μL of Nano-Glo Luciferase Substrate was added. The plate was incubated in the dark at room temperature for 3 min., and luminescence was measured using a Synergy HTX imaging multimode reader (BioTek, Winooski, USA).

#### Transfection of *C. parvum* sporozoites

Oocysts ($2.5\times10^7$ per transfection) were excysted as described above, and sporozoites released were resuspended in 80 μL complete SF buffer (consisting of 65.6 μL Cell Line Solution and 14.4 μL Supplement 1). Afterwards, 20 μL of plasmid suspension (50 μg Cas9/gRNA expression plasmid and 50 μg homologous repair plasmid) was added to the mixture, which was then transferred to a cuvette (Lonza Cologne, Cologne, Germany) and electroporated using an AMAXA 4D nucleofector system (Lonza Cologne) with program EH100. The transfected sporozoites were diluted in PBS and inoculated into three GKO mice by oral gavage. Prior to infection, 100 μL of 8% $NaHCO_3$ solution was administered to mice to neutralize gastric acid. On the next day, the drinking water in animal cages was replaced with a solution containing 16 g/L paromomycin (Yuaye, Shanghai, China) for *in vivo* selection. Parasite shedding was monitored by measuring nanoluciferase activity in the feces of infected mice.

#### Immunofluorescence microscopy assay

For IFA analysis of extracellular sporozoites and oocysts, oocysts were excysted and placed on a coverslip, air dried, fixed with 4% paraformaldehyde for 15 min, permeabilized with 0.5% Triton X-100 (Sigma-Aldrich, St. Louis, USA) at 37°C for 30 min, and blocked with 1% BSA at 37°C for 20 min. For IFA analysis of intracellular parasites *in vitro*, HCT-8 cells were infected with oocysts and incubated for different periods of time. The cells were fixed with 4% paraformaldehyde for 15 min, permeabilized with 0.1% Triton X-100 at 37°C for 15 min, and blocked with 1% BSA at 37°C for 20 min. The following primary antibodies were diluted in 0.1% BSA for staining: rabbit anti-HA (CST, Danvers, USA) at 1:800, mouse anti-FLAG (Beyotime, Shanghai, China) at 1:500, anti-GP40 (purified mouse mAbs) at 1:1000, anti-GP15 (purified mouse pAbs) w at 1:500, and anti-CP23 (rabbit pAb) at 1:1000. Cells were incubated with the antibodies overnight at 4°C, followed by 6 washes with PBS. Alexa Fluor-conjugated secondary antibodies (Thermo, Massachusetts, USA) diluted 1:1000 and Hoechst diluted 1:2000 in 0.1% BSA were added to the coverslips and incubated at 37°C for 30 min. Finally, the cells were washed 6 times in PBS, mounted on slides using Vectashield, and sealed with nail polish. The stained slides were examined under a Zeiss Axioskop Mot 2 fluorescence microscope (Carl Zeiss, Oberkochen, Germany).

**Immunoelectron microscopy (IEM)**
GKO mice were infected with 1000 transgenic oocysts and the ileum was harvested 12 days after infection. The samples were cut lengthwise, washed three times with cold PBS, and fixed at 4°C in freshly prepared 0.1% glutaraldehyde and 4% paraformaldehyde mix for 4 h. After gradient dehydration, infiltration in LR-White resin overnight and another 24 h in fresh LR-White resin at -20°C, the samples were transferred to fresh LR-White resin and polymerized under UV light (365 nm wavelength) at -25°C for 3 to 4 days. The samples were trimmed and cut into 70 nm ultrathin sections using a Leica EM UC7 ultramicrotome (Leica Microsystems Inc.). The sections were blocked with 1% BSA for 20 min and then incubated at 4°C with rabbit anti-HA antibody (CST) overnight. After washing in 0.1% BSA, the sections were further incubated with 10 nm colloidal gold-conjugated goat anti-rabbit IgG (H+L) for 60 min. Finally, the sections were stained with 2% uranyl acetate (w/v) and examined under a Talos L120C transmission electron microscope (Thermo). No IEM was done with the anti-FLAG mAb because of its low affinity, leading to the presence of few gold particles after sample processing.

**Scanning electron microscopy (SEM)**
The ileum was collected as described above, prefixed in 2.5% glutaraldehyde, postfixed in 0.2 M osmium tetroxide ($OsO_4$), and examined under an EVO MA 15/LS 15 scanning electron microscope (Carl Zeiss).

**Western blot analysis**
After excystation of $5\times10^6$ oocysts, sporozoites were lysed with 40 μL RIPA lysis buffer (Thermo) and incubated at 4°C overnight. The supernatant was collected after centrifugation and mixed with 10 μL 5×SDS loading buffer (TRANS, Beijing, China) before boiling for 10 min. Samples were stored at -20°C to prevent protein renaturation. The prepared samples were resolved by SDS-PAGE and transferred to a PVDF membrane. The membrane was then blocked in TBST (0.05% Tween 20 in TBS) with 5% skim milk at room temperature for 2 h, followed by three 5-min washes with TBST. The following primary antibodies were diluted in TBST containing 0.5% skim milk: rabbit anti-HA (CST) at 1:800, mouse anti-FLAG (Beyotime) at 1:400, and anti-NFDQ1 (purified rabbit pAbs) at 1:1000. The membrane was then incubated with the antibodies overnight at 4°C, followed by five washes of 5 min each with TBST. After overnight incubation at 4°C, the membrane was washed with TBST and then incubated with HRP-conjugated antibody (Beyotime) diluted 1:2000 in TBST containing 0.5% skim milk for 45 min at room temperature. After washing with PBST, the membrane was incubated with Immobilon Western Chemiluminescent HRP Substrate (1:1 solution A-solution B; Millipore), viewed on the Chemstudio imaging system (Analytik Jena, Germany), and imaged using VisionWorksLS (Analytik Jena, Germany).

**Primers**
PCR primers used in the study are listed in Supplementary Table 1 in the supplemental material. They were synthesized by Sangon Biotech (Shanghai, China).

**Construction of repair templates and CRISPR/Cas9 plasmids for genetic modification of *C. parvum***
Homology repair templates and CRISPR/Cas9 plasmids were generated as previously described (2). Briefly, CRISPR/Cas9 plasmids were generated by adding a single guide RNA (sgRNA) targeting the gene of interest to the linear Cas9 plasmid amplified from pACT:Cas9-GFP, U6:sgINS1 via Gibson assembly cloning. To target the *GP60* gene for depletion, a dual sgRNA strategy was used to target regions at the 5' and 3' ends of the gene. Primers for CRISPR/Cas9 are listed in Supplementary Table 1.
*Endogenous epitope tagging*
A dual sgRNA-directed CRISPR/Cas strategy was used for endogenous tagging of the *GP60* gene. The plasmid pACT1:Cas9-GFP, U6:sgGP60 was generated by replacing the INS1 sgRNA in the plasmid pACT1:Cas9-GFP, U6:sgINS1 (2) with an sgRNA corresponding to one region 43 bp downstream of the initiation codon and the other region 45 bp upstream of the stop codon in the *GP60* gene (CPCDC_6g1080) by Gibson assembly cloning using the ClonExpress II One Step Cloning Kit (Vazyme, Nanjing, China). The *GP60* sgRNA was designed using the eukaryotic

pathogen CRISPR guide RNA/DNA design tool (http://grna.ctegd.uga.edu). To generate a plasmid capable of tagging both the *GP40* fragment with the HA tag and the *GP15* fragment with the FLAG tag, a portion of the N-terminal UTR (412 bp), CDS (969 bp), and C-terminal UTR (888 bp) of the *GP60* gene was first amplified from *C. parvum* (IIdA20G1) genomic DNA. Then, the Nluc-P2A-neo reporter and the pUC19 backbone were amplified from the pINS1-3HA-Nluc-P2A-neo plasmid (2). After the plasmid pGP40/GP15-Nluc-P2A-neo was completed by Gibson assembly cloning, 3×HA and 3×FLAG were inserted into the region 99 bp downstream of the initiation codon and the other region 291 bp upstream of the stop codon in the *GP60* gene using the same approach. The tagging plasmid pGP40-HA-GP15-FLAG-Nluc-P2A-neo was completed after introduction of the mutant protospacer adjacent motif (PAM) using Gibson assembly cloning.

*Gene knockout*

To generate a plasmid for *GP60* gene depletion, the targeting plasmid p*GP60*-mCh-Nluc-P2A-neo-*GP60* was made by replacing the UPRT homologous flanks in plasmid UPRT-mCh-Nluc-P2A-neo-UPRT with *GP60* homologous flanks (1142 bp upstream and 1075 bp downstream of the CPCDC_6g1080 gene) using Gibson assembly of PCR-amplified fragments. To generate plasmids for *GP40* and *GP15* gene depletion, the targeting plasmids pmCh/GP15-Nluc-P2A-neo and pGP40/mCh-Nluc-P2A-neo were generated by modification of the plasmid pGP40/GP15-Nluc-P2A-neo described above. The mCherry sequence was amplified from UPRT-mCh-Nluc-P2A-neo-UPRT, and used to replace the *GP40* (*GP60* CDS 100~618 bp) or *GP15* (*GP60* CDS 676~870 bp) gene by Gibson assembly cloning. To knock out the furin-like protease cleavage sites (RSRR), signal peptide (SP) and glycosylphosphatidyl inositol (GPI) anchor in the *GP60* gene, the targeting plasmid pGP40Δ*RSRR*-HA/GP15-FLAG-Nluc-P2A-neo, pGP40Δ*SP*-HA/GP15-FLAG-Nluc-P2A-neo, and pGP40-HA/GP15Δ*GPI*-FLAG-Nluc-P2A-neo were prepared by modifications of the plasmid pGP40-HA/GP15-FLAG-Nluc-P2A-neo.

*Gene replacement*

To replace the *GP60* gene in the virulent IIdA20G1-HLJ isolate with the *GP60* sequence of the avirulent IIaA17G2R1-IOWA isolate, the targeting plasmid pGP40/GP15 (IIaA17G2R1)-Nluc-P2A-neo-mCh was made by modifying the plasmid pGP40/GP15 (IIdA20G1)-Nluc-P2A-neo. The *GP60*-IIdA20G1 sequence in the plasmid pGP40/GP15 (IIdA20G1)-Nluc-P2A-neo was replaced with the *GP60*-IIaA17G2R1 sequence amplified from DNA extracted from IIaA17G2R1-IOWA. The mCherry sequence and PAM were introduced into the plasmid as described above.

***In vitro* growth assay of *C. parvum***

HCT-8 cells were seeded in 48-well plates and cultured as described above, followed by infection with 10000 oocysts per well. Luminescence was measured at 3, 12, 24, 36, and 48 h after infection. The relative luminescence value was obtained by dividing the luminescence value of the corresponding day by the luminescence value at 3 h. The growth rate was calculated by dividing the following day's data by the previous day's data.

***In vivo* infected assay of *C. parvum***

IFN-γ KO mice or WT (C57BL/6) mice aged 3 to 5 weeks were used in the experiment. They were weighed on the day of infection, assigned to groups according to the experimental conditions, with the mean body weight being similar among groups. Mice were housed in individual cages during the entire infection studies. Each mouse was infected with 1000 fresh oocysts by gavage, and received paromomycin (16 g/L) in drinking water during the experiment. Feces were collected every two days, starting at DPI 2, and oocyst excretion intensity in feces was determined by OPG and luciferase assay. Mice were weighed every two days, and the time of death of mice was recorded for the preparation of the survival curve.

**Measurement of physical activity of mice**

A 45 s video was recorded for each control or infected mouse placed in clean cage. ImageJ was used to track the movement of mice in videos. The moving distance of the mouse within 45 s was calculated based on the movement trajectory, and the speed of movement was calculated using the result. GraphPad Prism 9.0 (GraphPad software) was used for data processing and statistical analysis.

**Quantification of oocyst shedding**
DNA was extracted from fecal pellets using a Magbeads FastDNA Kit for Soil (MP Biomedicals, Santa Ana, USA). Oocyst numbers were quantified by SSU rRNA-based qPCR as previously described (3). A standard curve for *C. parvum* genomic DNA was generated by extracting DNA from negative fecal samples spiked with 10, $10^2$, $10^3$, $10^4$, $10^5$, and $10^6$ oocysts of the *C. parvum* IOWA isolate (Waterborne, New Orleans, USA). The qPCR reactions were performed on a LightCycler 480 II (Roche, Indianapolis, USA) as previously described (4).

**PCR verification of gene integrations**
PCR analysis of the targeted genomic regions was used to confirm the proper integration of the insertion or replacement sequences in transgenic lines. The PCR reaction contained DNA purified from oocysts, 2×Fine Taq PCR SuperMix (TRANS, Beijing, China), and the primers listed in Supplementary Table 1. The PCR products generated were analyzed by electrophoresis and imaged on the GelDoc XR+ system (BioRad, Hercules, USA).

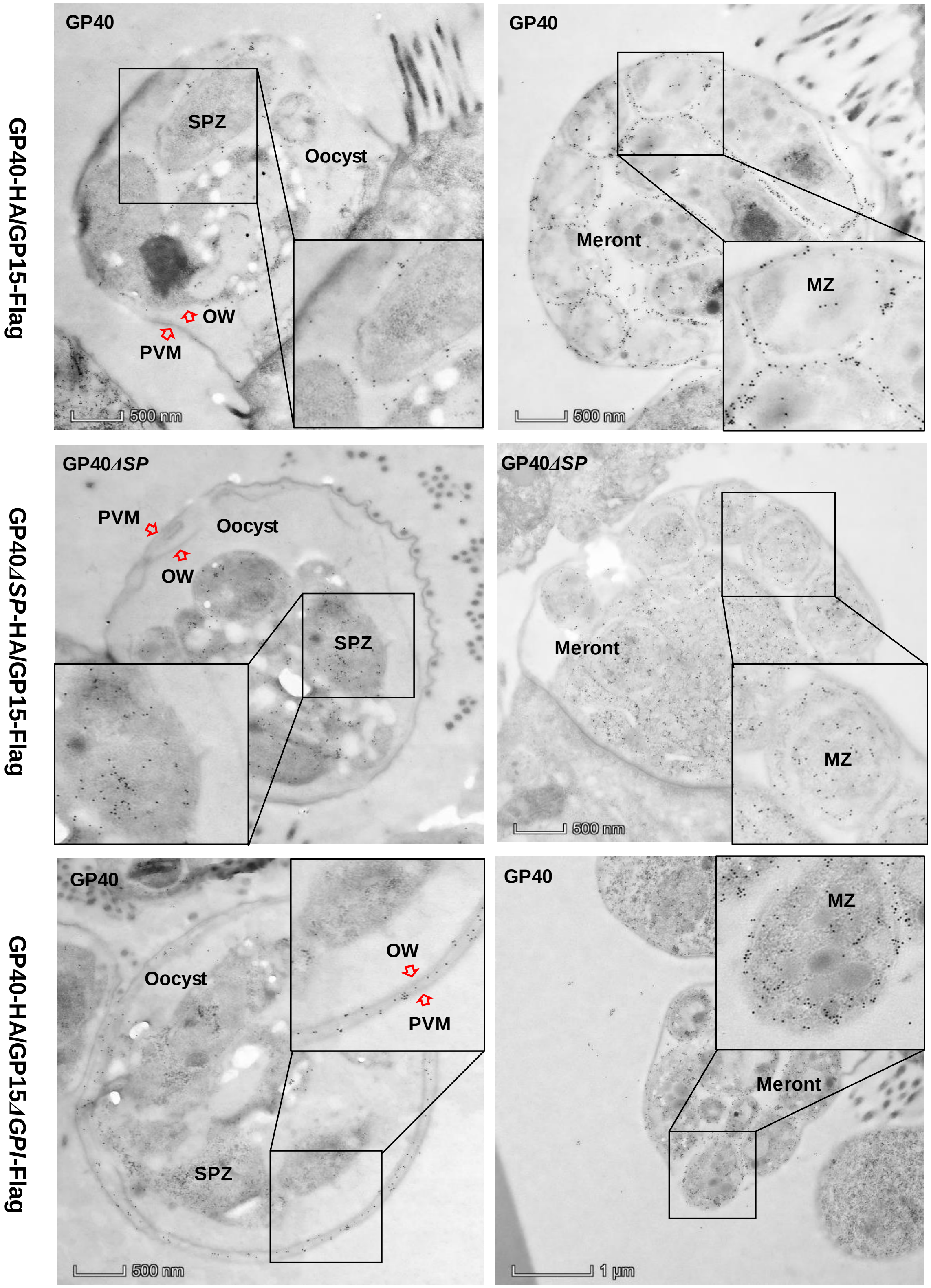


**Fig. S1.** Immunoelectron microscopy images of *Cryptosporidium parvum* oocysts and meronts in IFN-γ knockout (GKO) mice infected with the HA and FLAG-tagged WT line and similarly tagged but with the signal peptide (SP) or glycophosphatidylinositol (GPI) anchor depleted, using a mAb against recombinant GP40. OW: oocyst wall; PVM: parasitophorous vacuole membrane.

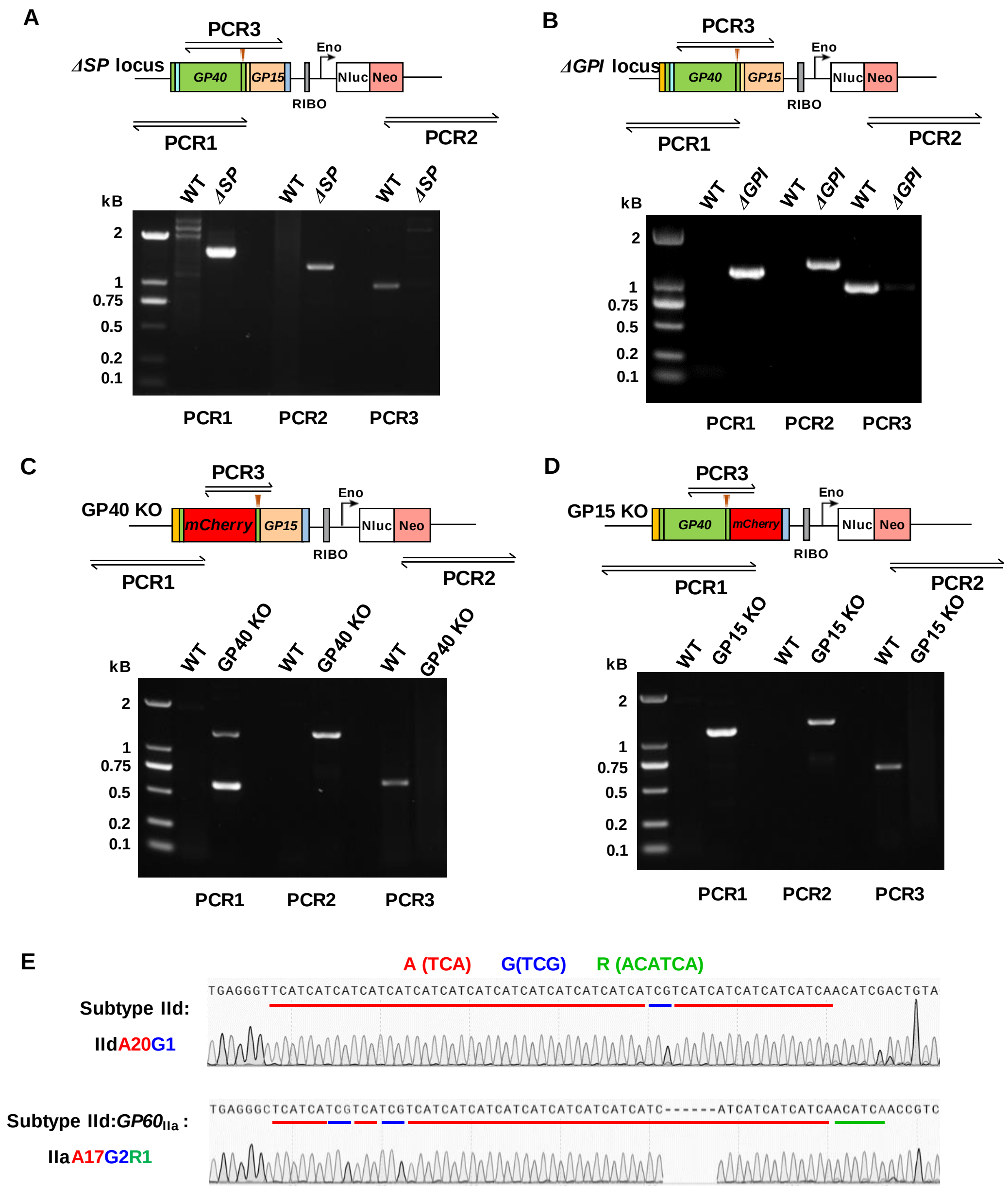


**Fig. S2.** Confirmation of the correct integration of the knockout and replacement cassette through PCR analysis. **A-D.** Results of PCR analysis of the gene integration. The design of PCR primers for the modified loci is shown above gel images of the PCR products. The gel images show the correct 5′ and 3′ integration of the repair DNA for all modified loci. PCR3 targets the native sequence of each locus and confirms the absence of WT parasites in the transgenic lines. **E.** Confirmation of the replacement of the GP60 nucleotide sequence by PCR and sequence analysis. Electropherogram of PCR products from HA- and FLAG-tagged WT line of the IIdA20G1 subtype (top) and replacement mutant of the IIaA17G2R1 subtype IIa (bottom). Subtype IId, the nucleotide sequence of the wild-type parasite. Subtype IId:$GP60_{IIa}$, the nucleotide sequence of the replacement mutant.

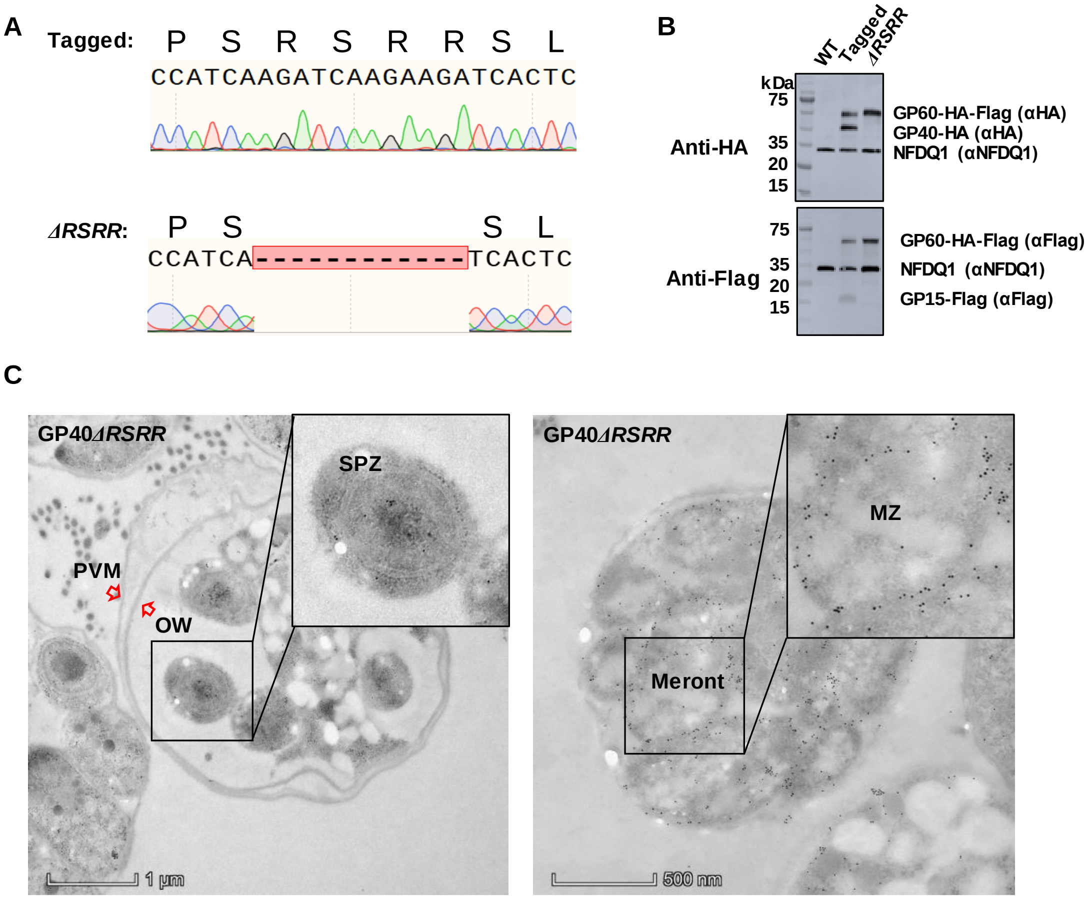


**Fig. S3.** Effects of depletion of the furin cleavage site on the processing of GP60 in *Cryptosporidium parvum*. **A.** Confirmation of the depletion of nucleotide sequence encoding furin cleavage sequence RSRR in the *GP60* gene by PCR and sequence analysis. **B.** Prevention of proteolytic cleavage of GP60 into GP40 (tagged with HA) and GP15 (tagged with FLAG) in RSRR-depleted line of *C. parvum* as indicated by Western blot analysis of protein extracted from oocysts using mAbs against the HA and FLAG tag. The HA and FLAG-tagged WT line is used as the control. **C.** Immunoelectron microscopy images of *C. parvum* oocysts and meronts in IFN-γ knockout (GKO) mice infected with the RSRR-depleted line, using the mAb against the HA-tag at the N-terminus of the GP40. GP40 is mostly expressed on the surface of sporozoites within the oocyst and merozoites within the meront.

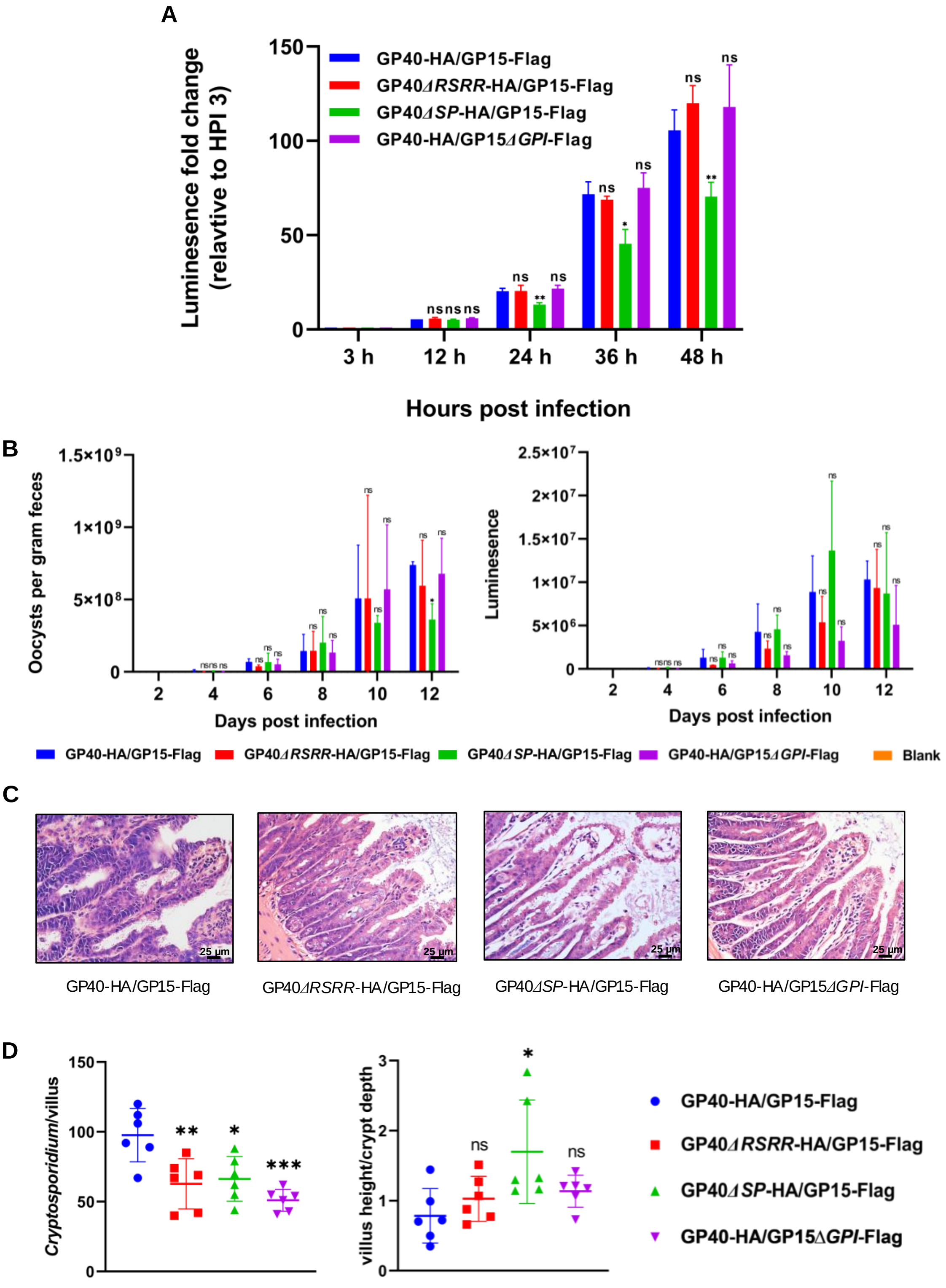


**Fig. S4.** Effects of depletion of the structural domains (signal peptide, GPI anchor, and furin cleavage site) in GP60 on *Cryptosporidium parvum* infection *in vitro* and *in vivo*. **A.** Effect of depletion of the signal peptide (SP), GPI anchor, and furin cleavage site (RSRR) in GP60 on the growth of *C. parvum* in HCT-8 cells in comparison with those infected with the HA and FLAG-tagged WT line. **B.** Effect of depletion of the structural domains on the infection intensity (as measured by oocysts per gram of feces and fecal luciferase activities) of *C. parvum*-infected IFN-γ knockout mice in comparison with those infected with the HA and FLAG-tagged WT line and the uninfected group. Mice infected with the RSRR- and GPI-depleted lines have better survival than those infected with the WT line. **C.** Comparison of the architecture of the ileal villi among mice infected with the transgenic lines above and the HA and FLAG-tagged WT line. **D.** Comparison of the villus height, crypt depth, and villus height to crypt depth ratio in the ileum among the groups above. Mice infected with the GPI-depleted line have significantly higher villus height to crypt depth ratio.

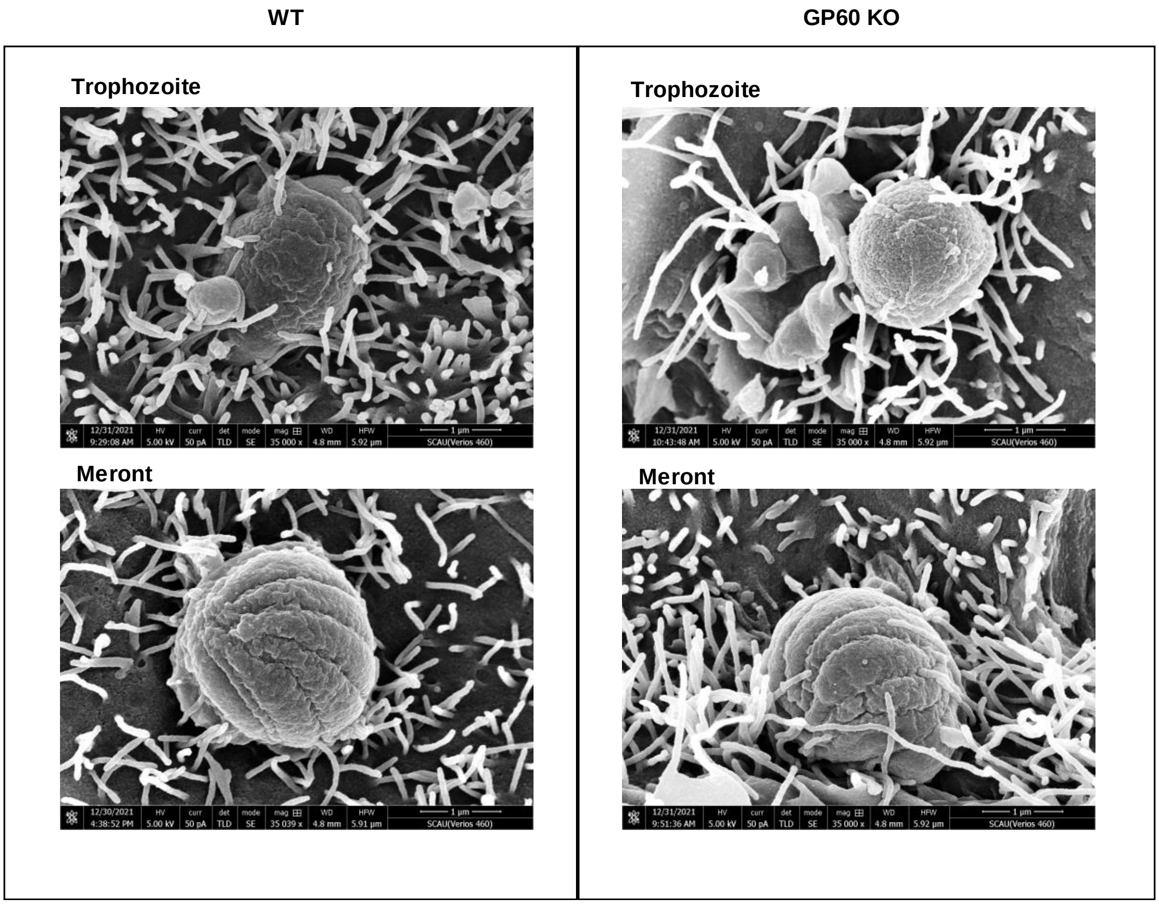


**Fig. S5.** Similarity in morphology of *Cryptosporidium parvum* stages between the GP60-depleted line and the HA and FLAG-tagged WT line, as indicated by scanning electron microscopy of infected HCT-8 cultures.

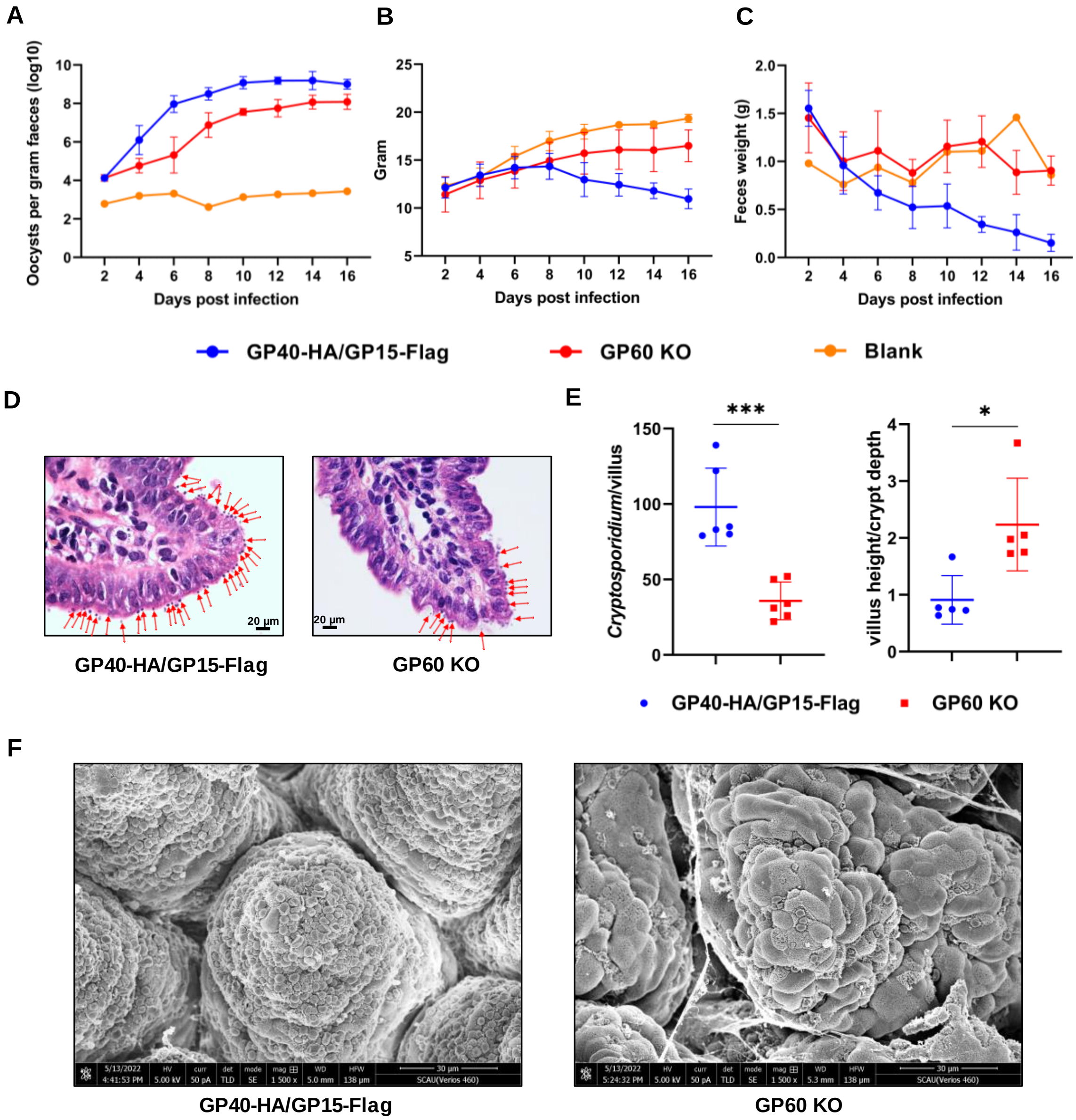


**Fig. S6.** Effects of GP60 depletion on the parasite load and virulence of *Cryptosporidium parvum*. **A-C**. Effect of GP60 depletion on the infection intensity (as measured by oocysts per gram of feces), body weight, and daily (a whole day) fecal weight of *C. parvum*-infected IFN-γ knockout mice in comparison with those infected with the HA and FLAG-tagged WT line and uninfected control group. **D & F**. Comparison of the parasite load (red arrows indicate parasites on the surface of the villus) on the surface of the ileal villi between mice infected with the GP60-depleted line above and the HA and FLAG-tagged WT line under H&E and scanning electron microscopy. The GP60 depletion has reduced the parasite burden significantly. **E**. Comparison of the villus height, crypt depth, and villus height to crypt depth ratio in the ileum between the two groups above. Mice infected with the GP60-depleted line have significantly higher villus height to crypt depth ratio.

**Tracks extracted from movies of behaving**

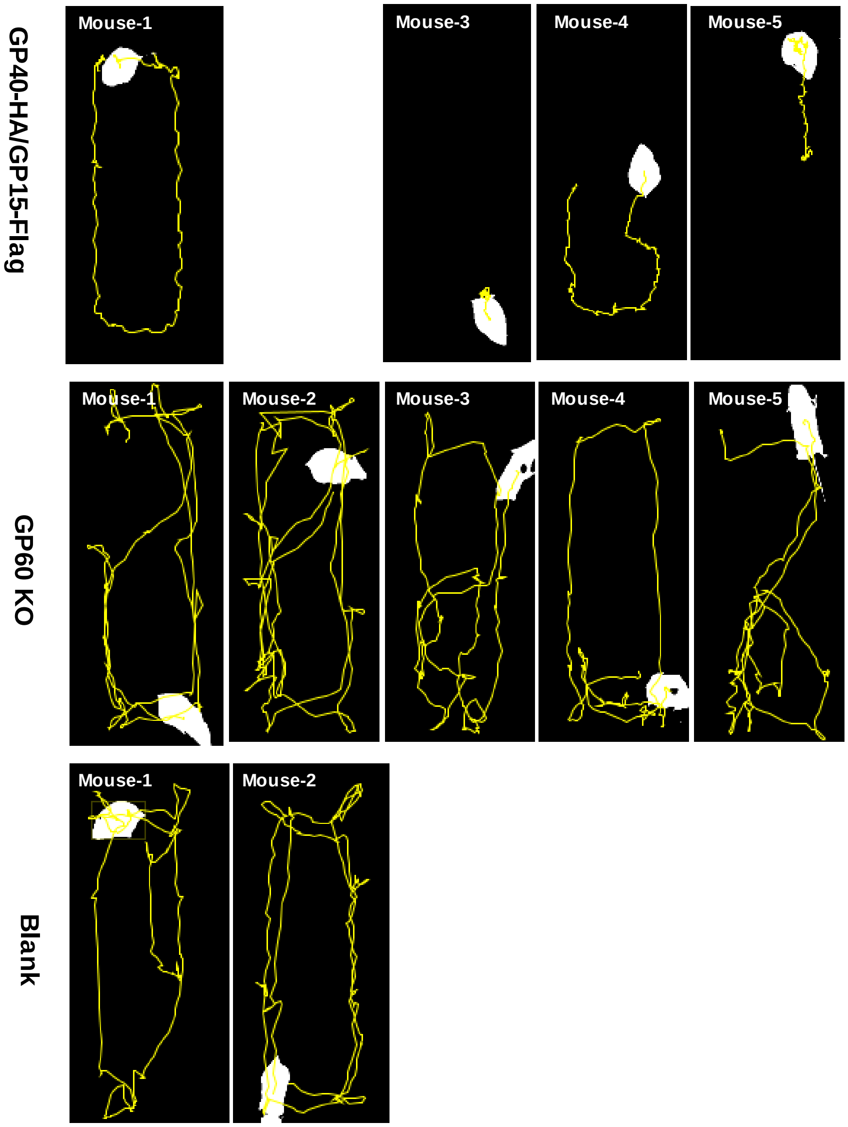


**Fig. S7.** Videos showing differences in physical movement between IFN-γ knockout mice infected with the GP60-depleted line (n = 5) and those infected with the HA and FLAG-tagged WT line (n = 5; one mouse died of the infection before the measurement), with the uninfected mice as controls (n = 2). Mice infected with the WT line show lethargy and arched back, while those infected with the GP60-depleted line show body posture and movements similar to the uninfected control mice.

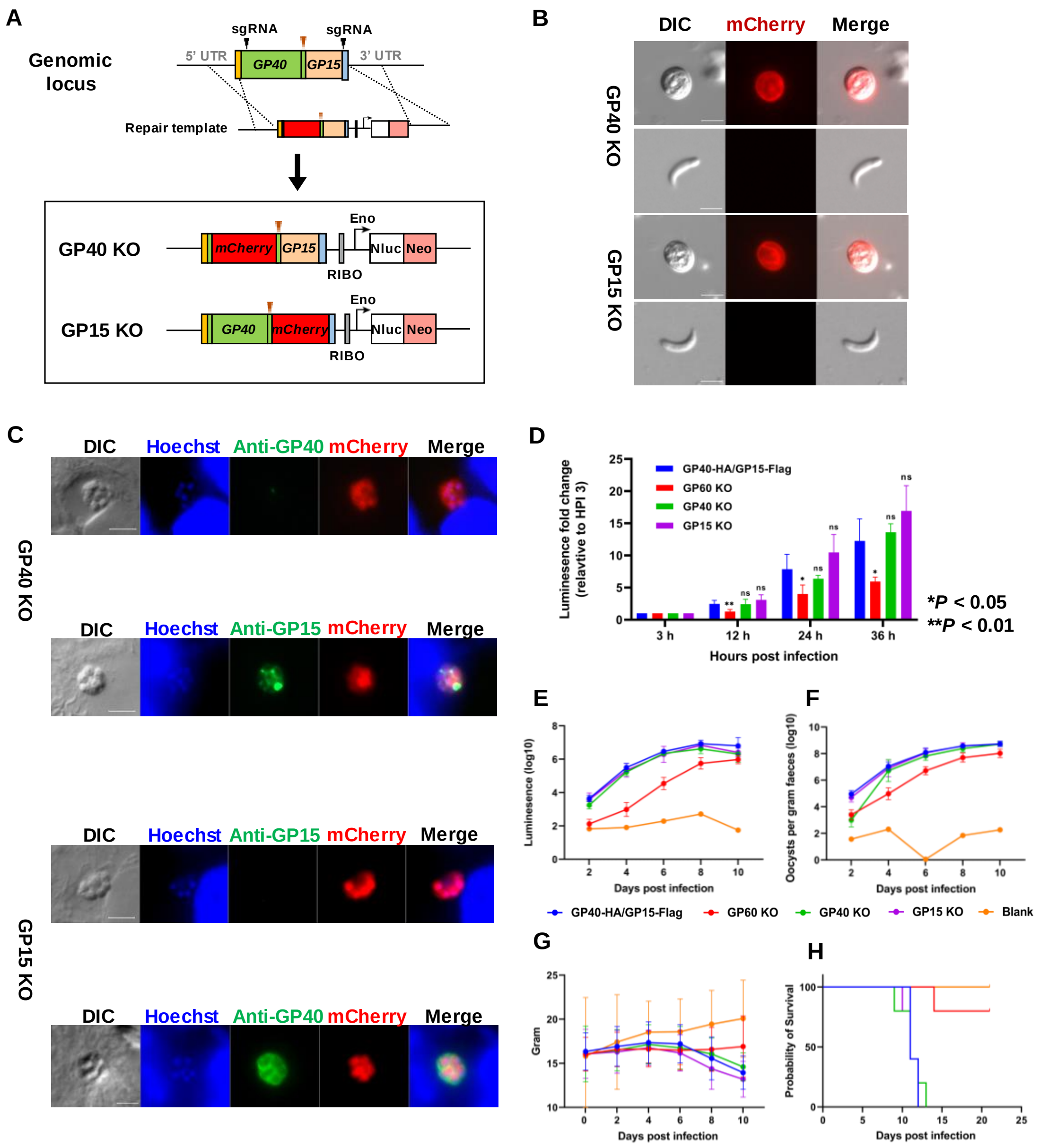


**Fig. S8.** Depletion of GP40 and GP15 individually have less effects on growth of *Cryptosporidium parvum*. **A.** Strategy used in the knockout of the *GP40* and *GP15* fragments of the *GP60* gene. **B.** Microscopy of oocysts from mice infected with the GP40 and GP15 knockout lines, with the transgenic oocysts being red and free sporozoites being colorless. **C.** Confirmation of the depletion of GP40 and GP15 through immunofluorescence analysis using a mAb against GP40 and polyclonal antibody against GP15. No GP40 is expressed on mCherry-positive parasites in HCT-8 cultures infected with the GP40-knockout line, while no GP15 is expressed on mCherry-positive parasites in HCT-8 cultures infected with the GP15-knockout line. **D.** Effect of GP40 and GP15 depletion on the growth of *C. parvum* in HCT-8 cells in comparison with those infected with the HA and FLAG-tagged WT line, the GP60-knockout line, and the uninfected control. **E-H.** Depletion of GP40 and GP15 have no significant effects on growth of *Cryptosporidium parvum* in INF-γ knockout mice. Effect of GP40 and GP15 depletion on the infection intensity of *C. parvum* in IFN-γ knockout mice as measured by fecal luciferase activity and oocysts per gram of feces, and bodyweight and survival of infected mice, in comparison with the three control groups above.

Only the GP60 depletion has reduced the infection intensity and improved the bodyweight and survival in *C. parvum*-infected mice.

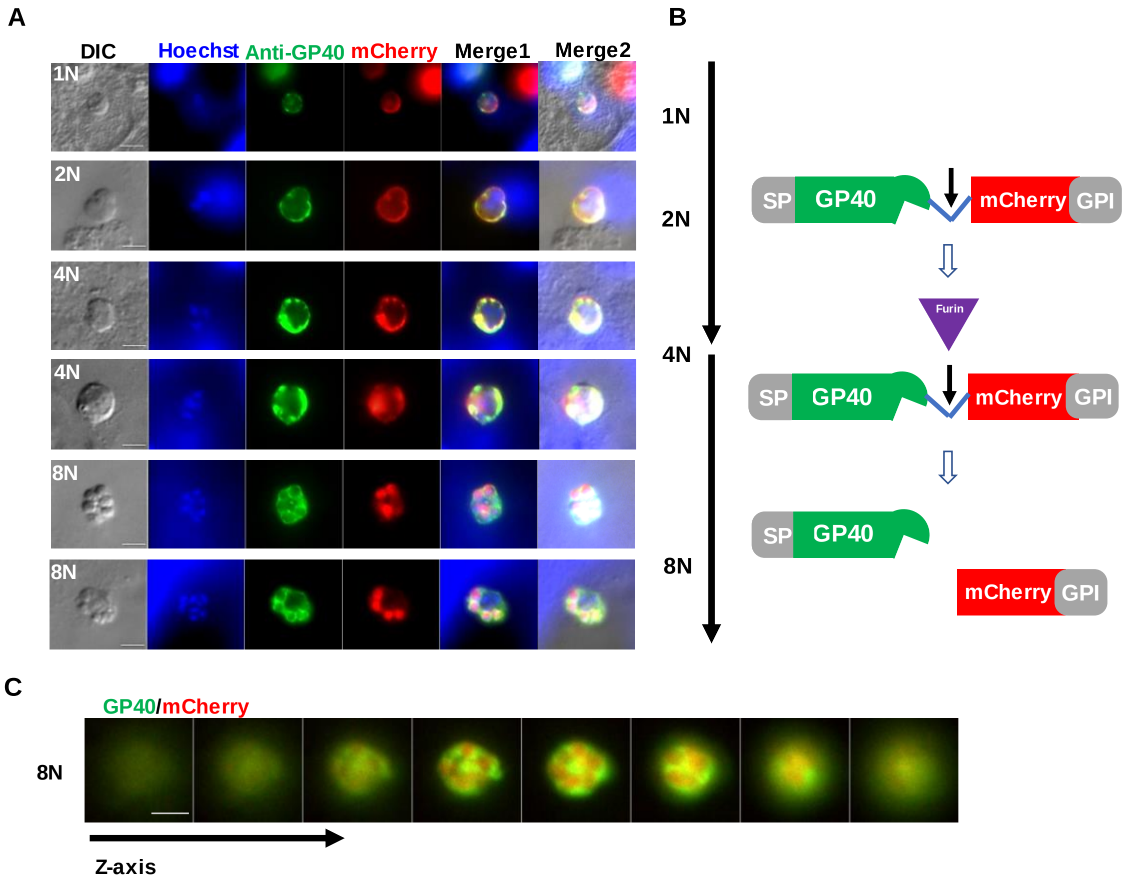


**Fig. S9.** GP15 depletion changed the location of GP40 in mature meronts. **A.** Immunofluorescence analysis of *Cryptosporidium parvum* trophozoites and meronts cultured in HCT-8 cells with the GP15 deletion, using a mAb against recombinant GP40; **B.** Schematic of the separation of GP40 and mCherry; **C.** Longitudinal observation of GP40 and mCherry localization in GP15 depletion mature meronts. GP40 and the mCherry did not co-localize with each other, with mCherry remaining inside merozoites.

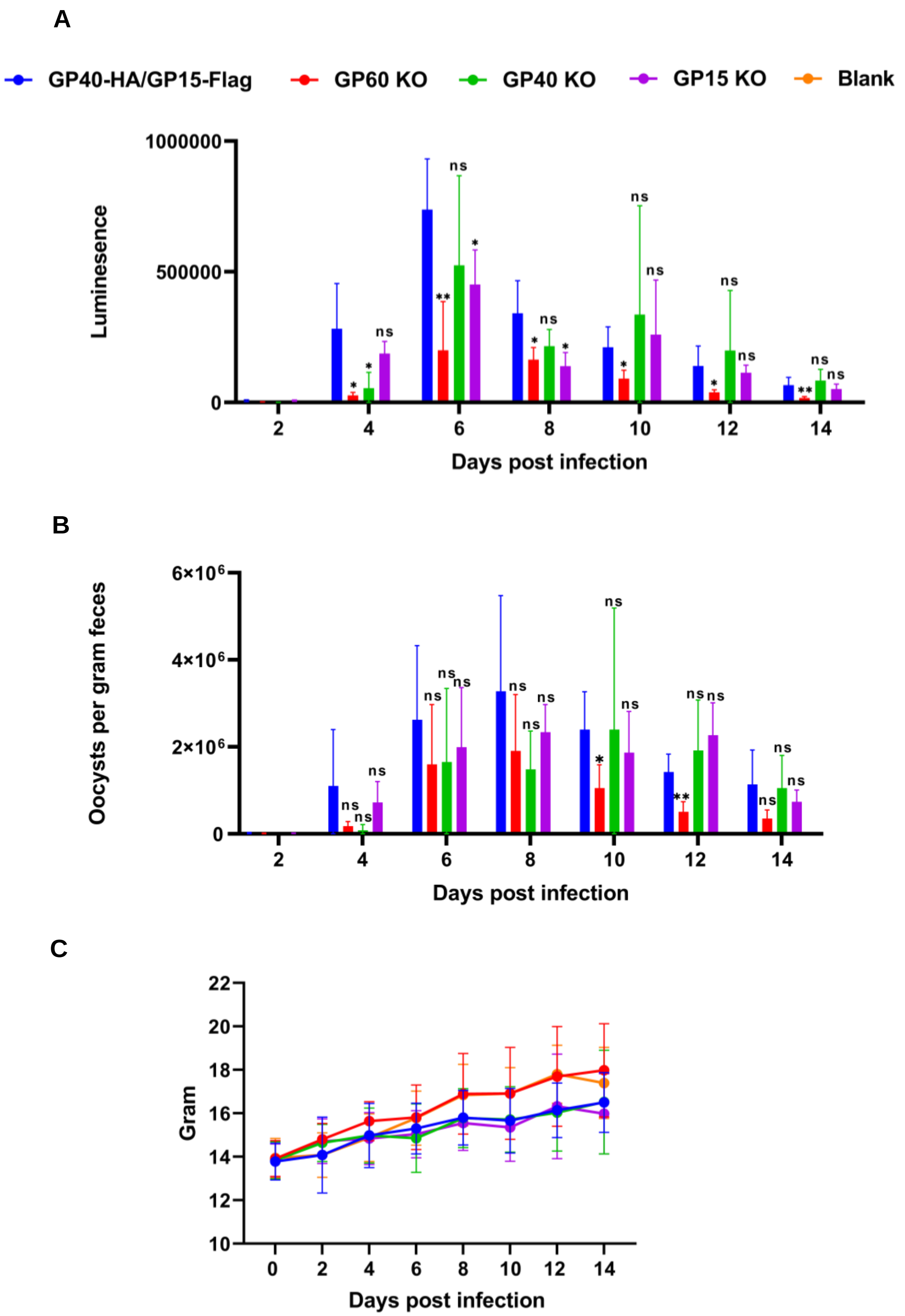


**Fig. S10.** Depletion of GP40 and GP15 have significant effects on growth of *Cryptosporidium parvum* in WT C57BL/6 mice. **A-C.** Effect of GP40 and GP15 depletion on the infection intensity of *C. parvum* in WT mice (C57BL/6) as measured by fecal luciferase activity (*A*) and oocysts per gram of feces (*B*), and bodyweight (*C*) in comparison with the GP60-depleted and the HA and

FLAG-tagged WT lines. GP60 depletion reduced the infection intensity and improved the bodyweight, while GP40 or GP15 depletion only reduced the in vivo infection intensity modestly.

**Table S1.** Primers used in this study.

| Primer | Primer Sequence | Use |
| --- | --- | --- |
| GP60clone-F | 5'-CGGCTCGACCCTTCTATAGGTG-3' | forward primer for cloning full-length GP60 |
| GP60clone-R | 5'-ACTGGCGTCCTGATTCAATG-3' | reverse primer for cloning full-length GP60 |
| GP60KO-gRNA1-F | 5'- CTATTTCTAGCTCTAAAACCTGCTTTTGGACTCAGATACCCCAACACTTAACCTTTC-3' | forward primer for amplifying GP60 gRNA1 fragment for GP60 knock out CRISPR plasmid construction |
| GP60KO-gRNA1-R | 5'-GAAAGGTTAAGTGTTGGGGTATCTGAGTCCAAAAGCAGGTTTTAGAGCTAGAAATAG-3' | reverse primer for amplifying GP60 gRNA1 fragment for GP60 knock out CRISPR plasmid construction |
| GP60KO-gRNA2-F | 5'-CTATTTCTAGCTCTAAAACCTGCTTTTGGACTCAGATACCCCAACACTTAACCTTTC-3' | forward primer for amplifying GP60 gRNA2 fragment for GP60 knock out CRISPR plasmid construction |
| GP60KO-gRNA2-R | 5'-GAAAGGTTAAGTGTTGGGGTATCTGAGTCCAAAAGCAGGTTTTAGAGCTAGAAATAG-3' | reverse primer for amplifying GP60 gRNA2 fragment for GP60 knock out CRISPR plasmid construction |
| GP60KOcassette-F | 5'-CTATGCAGGCATGCAAGCTTCCAGTGAATTCGAGCGCC-3' | forward primer for amplifying cassette containing GP60 gRNA1 fragment for dual sgRNA CRISPR plasmid construction |
| GP60KOcassette-R | 5'-ACAGGAAACAGCTATGACCCAAATTTCTCCAACCAGT-3' | reverse primer for amplifying cassette containing GP60 gRNA1 fragment for dual sgRNA CRISPR plasmid construction |
| GP60KOvector-F | 5'-GTCATAGCTGTTTCCTGTGTGAAATTG-3' | forward primer for amplifying vector for dual sgRNA CRISPR plasmid construction |
| GP60KOvector-R | 5'-GCTTGCATGCCTGCATAGTATATCC-3' | reverse primer for amplifying vector for dual sgRNA CRISPR plasmid construction |
| pActin-mCherry-TerActin-nluc-neo-F | 5'-AGGCAACTAAGGACAAAGGAAGTCAGAATGAGTTGGTTATAAACAG-3' | forward primer for amplifying Actin-mCherry-TerActin-nluc-neo fragment for GP60 knock out plasmid construction |
| pActin-mCherry-TerActin-nluc-neo-R | 5'-ATCCGCGTGACAGGCGGATATACTAGGTGGGGAAACTAAATATAC-3' | reverse primer for amplifying Actin-mCherry-TerActin-nluc-neo fragment for GP60 knock out plasmid construction |
| GP60KOdonor5H-F | 5'-AGTCACGACGTTGTAAAACGTGCTAACAAAAGTGAATACAAC-3' | forward primer for amplifying GP60 5'UTR for GP60 knock out plasmid construction |
| GP60KOdonor5H-R | 5'-CTTCCTTTGTCCTTAGTTG-3' | reverse primer for amplifying GP60 5'UTR for GP60 knock out plasmid construction |
| GP60KOdonor3H-F | 5'-GCATGCAATCTATAACTAATTC-3' | forward primer for amplifying GP60 3'UTR for GP60 knock out plasmid construction |
| GP60KOdonor3H-R | 5'-TCTTAAAACTCTTAAACTTTATTATACTTCCTTGCTGGTTG-3' | reverse primer for amplifying GP60 3'UTR for GP60 knock out plasmid construction |
| Gp60KO-Vector-F | 5'-AAAGTTTAAGAGTTTTAAGAG-3' | forward primer for amplifying pUC19 vector for GP60 knock out plasmid construction |
| Gp60KO-Vector-R | 5'-CTTCCTTTGTCCTTAGTTGC-3' | reverse primer for amplifying pUC19 vector for GP60 knock out plasmid construction |

| Primer | Primer Sequence | Use |
|---|---|---|
| GP60KOPCR1-F | 5'-ATTGATCTGCAGTAATGTCTTAATG-3' | PCR1 of GP60 mutant (GP60 KO, GP40 KO, GP15 KO) |
| GP60KOPCR1-R | 5'-CCAACTCATTCTGACTTCC-3' | PCR1 of GP60 mutant (GP60 KO, GP40 KO, GP15 KO) |
| GP60KOPCR2-F | 5'-CAACGTATCGCCTTCTATCGTC-3' | PCR2 of GP60 mutant (GP60 KO, GP40 KO, GP15 KO) |
| GP60KOPCR2-R | 5'-TGAGCAAGAAGAGACACTCG-3' | PCR2 of GP60 mutant (GP60 KO, GP40 KO, GP15 KO) |
| GP60KOPCR3-F | 5'-ATGAGATTGTCGCTCATTATCG-3' | PCR3 of GP60 mutant (GP60 KO, GP40 KO, GP15 KO) |
| GP60KOPCR3-R | 5'-ACACGAATAAGGCTGCAAAG-3' | PCR3 of GP60 mutant (GP60 KO, GP40 KO, GP15 KO) |
| GP40KO-mCherry-F | 5'-GCACTTTAAAGGATGTTTCTGCCATCATCAAGGAGTTCATG-3' | forward primer for amplifying mCherry for GP40 knock out plasmid construction |
| GP40KO-mCherry-R | 5'-TTCTTGATCTTGATGGAACCTGACC CTTGTACAGCTCGTCCATGC-3' | reverse primer for amplifying mCherry for GP40 knock out plasmid construction |
| GP40KO-vector-F | 5'-GGTCAGGTTCCATCAAGATCAAG-3' | forward primer for amplifying vector for GP40 knock out plasmid construction |
| GP40KO-vector-R | 5'-GATCTTGATGGAACCTGACC-3' | reverse primer for amplifying vector for GP40 knock out plasmid construction |
| GP15KO-mCherry-F | 5'-TCACTCTCAGAGGAGGCTAGTGAAACTGCAGCCATCATCAAGGAGTTCATG-3' | forward primer for amplifying mCherry for GP15 knock out plasmid construction |
| GP15KO-mCherry-R | 5'-TCCTTCAAAAGAACTTTGTT CTTGTACAGCTCGTCCATGC-3' | reverse primer for amplifying mCherry for GP15 knock out plasmid construction |
| GP15KO-vector-F | 5'-AACAAAGTTCTTTTGAAGGATG-3' | forward primer for amplifying vector for GP15 knock out plasmid construction |
| GP15KO-vector-R | 5'-TGCAGTTTCACTAGCCTCCTC-3' | reverse primer for amplifying vector for GP15 knock out plasmid construction |
| GP60-tagging-F | 5'-GGGTTTTCCCAGTCACGACGTTGTAAAACGCCAATATTTGTGCATTATACG-3' | forward primer for amplifying GP60 for GP40/GP15-Nluc-P2A-neo donor plasmid construction |
| GP60-tagging-R | 5'-CCTGATTCAATGTAATAAACAAAACCTGATAAACGAGTACTTATA-3' | reverse primer for amplifying GP60 for GP40/GP15-Nluc-P2A-neo donor plasmid construction |
| GP40/GP15-Nluc-P2A-neo-Vector1-F | 5'-GGCGTAATCATGGTCATAGCTG-3' | forward primer for amplifying vector1 for GP40/GP15-Nluc-P2A-neo donor plasmid construction |
| GP40/GP15-Nluc-P2A-neo-Vector1-R | 5'-CGTTTTACAACGTCGTGACTG-3' | reverse primer for amplifying vector1 for GP40/GP15-Nluc-P2A-neo donor plasmid construction |
| RIBO-nluc-neo-F | 5'-TGCAGCCTTATTCGTGTTGTAA TTAATTAAGAAATCTTTTTTAGCTGG-3' | forward primer for amplifying RIBO-nluc-neo for GP40/GP15-Nluc-P2A-neo donor plasmid construction |
| RIBO-nluc-neo-R | 5'-TTCAATGTAATAAACAAAAC TCAGAAGAATTCGTCAAGAAG-3' | reverse primer for amplifying RIBO-nluc-neo for GP40/GP15-Nluc-P2A-neo donor plasmid construction |
| GP40/GP15-Nluc-P2A-neo-Vector2-F | 5'-GTTTTGTTTATTACATTGAATC-3' | forward primer for amplifying vector2 for GP40/GP15-Nluc-P2A-neo donor plasmid construction |

| Primer | Primer Sequence | Use |
|---|---|---|
| GP40/GP15-Nluc-P2A-neo-Vector2-R | 5'-TTACAACACGAATAAGGCTGC-3' | reverse primer for amplifying vector2 for GP40/GP15-Nluc-P2A-neo donor plasmid construction |
| mutantPAM1-F | 5'-TTACTCTCCGTTATAGTCTCGGCTGTATTCTCAGCCCCAGCCGTTC-3' | forward primer for mutating PAM of gRNA1 sequence |
| mutantPAM1-R | 5'-ACAGCCGAGACTATAACGGAG-3' | reverse primer for mutating PAM of gRNA1 sequence |
| mutantPAM2-F | 5'-GAAGGATGCTGGTTCGTCTGCTTTTGGACTCAGATACATCGTTC-3' | forward primer for mutating PAM of gRNA2 sequence |
| mutantPAM2-R | 5'-AAGCAGACGAACCAGCATCC-3' | reverse primer for mutating PAM of gRNA2 sequence |
| GP40-HA/GP15-Nluc-P2A-neo-vector-F | 5'-GTTGAGGGTTCATCATCATC-3' | forward primer for amplifying vector for GP40-HA/GP15-Nluc-P2A-neo donor plasmid construction |
| GP40-HA/GP15-Nluc-P2A-neo-vector-R | 5'-AGAAACATCCTTTAAAGTGCCTC-3' | reverse primer for amplifying vector forGP40-HA/GP15-Nluc-P2A-neo donor plasmid construction |
| 3×HA-F | 5'-TCAGAGGCACTTTAAAGGATGTTTCT TACCCATACGATGTTCCAG-3' | forward primer for amplifying 3×HA for GP40-HA/GP15-Nluc-P2A-neo donor plasmid construction |
| 3×HA-R | 5'-ATGATGATGATGATGAACCCTCAAC AGCGTAATCTGGAACGTCATATG-3' | reverse primer for amplifying 3×HA forGP40-HA/GP15-Nluc-P2A-neo donor plasmid construction |
| 3×Flag-F | 5'-GATCATGATATTGATTACAAAGACGATGACGATAAG ACCGTCGATTTGTTTGCCTTCAC-3' | forward primer for GP40-HA/GP15-Flag-Nluc-P2A-neo donor plasmid construction |
| 3×Flag-R | 5'-CGTCTTTGTAATCAATATCATGATCCTTGTAGTCTCCGTCGTGGTCCTTATAGTC TGCAGTTTCACTAGCCTCCTC-3' | reverse primer for GP40-HA/GP15-Flag-Nluc-P2A-neo donor plasmid construction |
| GP40-HA/GP15-Flag PCR1 identify-F | 5'-CGTAGGCTTGAGTGCTCAAC-3' | PCR1 of GP60 mutant (GP60 Tagging, GP60 SP KO, GP60 GPI KO, GP60 RSRR KO) |
| GP40-HA/GP15-Flag PCR1 identify-R | 5'-ATTGAAGCAGATCTCGGCAGAC-3' | PCR1 of GP60 mutant (GP60 Tagging, GP60 SP KO, GP60 GPI KO, GP60 RSRR KO) |
| GP40-HA/GP15-Flag PCR2 identify-F | 5'-CAACGTATCGCCTTCTATCGTC-3' | PCR2 of GP60 mutant (GP60 Tagging, GP60 SP KO, GP60 GPI KO, GP60 RSRR KO) |
| GP40-HA/GP15-Flag PCR2 identify-R | 5'-TGAGCAAGAAGAGACACTCG-3' | PCR2 of GP60 mutant (GP60 Tagging, GP60 SP KO, GP60 GPI KO, GP60 RSRR KO) |
| Subtype IId-Vector-F | 5'-GTTTATTACATTGAATCAGGACG-3' | forward primer for amplifying vector for Subtype IId donor plasmid construction |
| Subtype IId-Vector-R | 5'-GAAGAATTCGTCAAGAAGAC-3' | reverse primer for amplifying vector for Subtype IId donor plasmid construction |
| P2A-F | 5'-TTCTATCGTCTTCTTGACGAATTCTTC GGAAGCGGAGCTACTAACTTC-3' | forward primer for P2A fragment for Subtype IId donor plasmid construction |

| Primer | Primer Sequence | Use |
|---|---|---|
| P2A-R | 5'-GAAGCGCATGAACTCCTTGATGATGGCCAT AGGTCCAGGGTTCTCCTCCAC-3' | reverse primer for P2A fragment for Subtype IId donor plasmid construction |
| Subtype IId:GP60IIa-mCherry-F | 5'-ATGGCCATCATCAAGGAGTTC-3' | forward primer for mCherry fragment for Subtype IId donor plasmid construction |
| Subtype IId:GP60IIa-mCherry-R | 5'-AAACTGGCGTCCTGATTCAATGTAATAAAC TTACTTGTACAGCTCGTCCATG-3' | reverse primer for mCherry fragment for Subtype IId donor plasmid construction |
| IIa-GP60-F | 5'-AGTCTCGGCTGTATTCTCAGCC CCAGCCGTTCCACTCAGAGG-3' | forward primer for IIa GP60 for Subtype IId:$GP60_{IIa}$ donor plasmid construction |
| IIa-GP60-R | 5'-AGATTGCAAAAACGGAAGGAACGATGTATCTGAGTCCAAAAGCAGAC-3' | reverse primer for IIa GP60 fragment for Subtype IId:$GP60_{IIa}$ donor plasmid construction |
| Subtype IId:$GP60_{IIa}$-Vector-F | 5'-ATCGTTCCTTCCGTTTTTGC-3' | forward primer for vector for Subtype IId:$GP60_{IIa}$ donor plasmid construction |
| Subtype IId:$GP60_{IIa}$-Vector-R | 5'-GGCTGAGAATACAGCCGAGAC-3' | reverse primer for vector forSubtype IId:$GP60_{IIa}$ donor plasmid construction |
| GP40*Δ*SP-HA/GP15-Flag-F | 5'-TAAGTAGGCAACTAAGGACAAAGGAAGATGCCAGCCGTTCCACTCAGAG-3' | forward primer for amplifying donor plasmid for GP60 SP knock out plasmid construction |
| GP40*Δ*SP-HA/GP15-Flag-R | 5'-CAACTAAGGACAAAGGAAGATG-3' | reverse primer for amplifying donor plasmid for GP60 SP knock out plasmid construction |
| GP40-HA/GP15*Δ*GPI-Flag-F | 5'-AATGAGGACGGAGACTTGGTTGACAAGGACTAA TTAATTAAGAAATCTTTTTTAG-3' | forward primer for amplifying donor plasmid for GP60 GPI knock out plasmid construction |
| GP40-HA/GP15*Δ*GPI-FlagR | 5'-GTCCTTGTCAACCAAGTCTCCGTC-3' | reverse primer for amplifying donor plasmid for GP60 GPI knock out plasmid construction |
| GP40*Δ*RSRR-HA/GP15-Flag-F | 5'-CGGATCTGCAGGTCAGGTTCCATCA TCACTCTCAGAGGAGGCTAGTG-3' | forward primer for amplifying donor plasmid for GP60 RSRR knock out plasmid construction |
| GP40*Δ*RSRR-HA/GP15-Flag-R | 5'-TGATGGAACCTGACCTGCAG-3' | reverse primer for amplifying donor plasmid for GP60 RSRR knock out plasmid construction |
